# Dose-dependent sensitivity of human 3D chromatin to a heart disease-linked transcription factor

**DOI:** 10.1101/2025.01.09.632202

**Authors:** Zoe L. Grant, Shuzhen Kuang, Shu Zhang, Abraham J. Horrillo, Kavitha S. Rao, Vasumathi Kameswaran, Carine Joubran, Pik Ki Lau, Keyi Dong, Bing Yang, Weronika M. Bartosik, Nathan R. Zemke, Bing Ren, Irfan S. Kathiriya, Katherine S. Pollard, Benoit G. Bruneau

**Author notes:** These authors contributed equally (Zoe L. Grant, Shuzhen Kuang).

## Abstract

Dosage-sensitive transcription factors (TFs) underlie altered gene regulation in human developmental disorders, and cell-type specific gene regulation is linked to the reorganization of 3D chromatin during cellular differentiation. Here, we show dose-dependent regulation of chromatin organization by the congenital heart disease (CHD)- linked, lineage-restricted TF TBX5 in human cardiomyocyte differentiation. Genome organization, including compartments, topologically associated domains, and chromatin loops, are sensitive to reduced *TBX5* dosage in a human model of CHD, with variations in response across individual cells. Regions normally bound by TBX5 are especially sensitive, while co-occupancy with CTCF partially protects TBX5-bound TAD boundaries and loop anchors. These results highlight the importance of lineage-restricted TF dosage in cell-type specific 3D chromatin dynamics, suggesting a new mechanism for TF-dependent disease.

## Introduction

Our understanding of the genetic foundation of most human developmental disorders primarily stems from dominant mutations in gene regulators, notably transcription factors (TFs) and chromatin-modifying factors (*1, 2*). Many of these mutations are predicted or shown to result in haploinsufficiency, where loss of a single copy of a gene leads to disease. The TFs involved are well-studied in the context of embryonic development. For example, PAX6 in aniridia, SOX9 in campomelic dysplasia, NOTCH1 in bicuspid aortic valve and TBX5, NKX2-5 and GATA4 in congenital heart defects (CHDs) (*2*). The major consequence of TF haploinsufficiency is transcriptional dysregulation. However, how TF haploinsufficiency regulates downstream gene regulatory networks is not mechanistically understood, despite decades of study.

The 3D genome is organized into hierarchical layers that confer regulation of transcriptional activity through formation of active (A) and repressive (B) compartments, topologically associating domains (TADs) and chromatin loops (*3*). The formation of three-dimensional, long-range interactions between cis-regulatory elements such as enhancers and promoters is especially important for regulating cell type specific gene expression (*4, 5*). CTCF binding in convergent orientation is often found at loop anchors, which acts as a docking site for cohesin complexes (*6–8*) following cohesin-dependent loop extrusion (*9, 10*). Different cell types are characterized by lineage-specific A/B compartments, TADs and chromatin loops (*11, 12*). Since CTCF and cohesin complex members are widely expressed across cell types, additional mechanisms must be responsible for cell-type-specific 3D genome organization. This includes lineage-constrained TFs as direct regulators of 3D genome organization (*13–18*). Given the specificity of TF expression, it is likely that a unique TF or set of TFs regulates 3D genome organization in each cell type.

The T-box TF TBX5 is a master regulator of cardiac gene expression required for normal heart patterning and morphogenesis (*19*). TBX5 dose-dependently regulates target gene expression (*19–21*) and the relevance of this is underscored in human disease, where heterozygous loss of function mutations in *TBX5* lead to Holt-Oram Syndrome (*22, 23*). These haploinsufficient mutations cause 85% penetrant CHDs, that are primarily atrial and ventricular septal defects and conduction system defects (*22, 23*). Using an induced pluripotent stem cell (iPSC) *TBX5* allelic series comprising wildtype (WT) and heterozygous or homozygous loss of function mutations we previously characterized the dosage-sensitive transcriptional changes and impact on gene-regulatory networks in cardiomyocytes (CMs) (*20*). Whether these TBX5 CHD-causing mutations directly impact 3D chromatin structures or influence generally how 3D chromatin is established remains to be determined.

Here, we sought to understand lineage-specific 3D chromatin regulation in the context of human CM differentiation by studying the CHD-relevant, lineage-restricted TF TBX5. We found extensive, dynamic reorganization of the 3D genome across atrial and ventricular CM differentiation with clear distinctions between the two CM types, in both bulk populations and at the single cell level. We observed that during cardiac differentiation TBX5 binding is enriched at cardiac lineage specific TAD boundaries and loop anchors. Importantly, our work identified that in the context of haploinsufficiency, TBX5 directly influences 3D genome organization including higher order organization of chromosomal compartments, TADs, and loops. These phenotypes were exacerbated by complete loss of TBX5, highlighting the key dosage-sensitive regions of the genome.

Overall, our results demonstrate that cell-type specific 3D genome organization is sensitive to the dosage of a lineage-restricted TF and suggest a novel mechanism by which reduced TF dosage may contribute to CHDs.

## Results

### Dynamic reorganization of the 3D genome during human cardiac differentiation

Analyses of 3D chromatin during CM differentiation have been carried out before (*24, 25*) but not at the resolution that would enable precise detection of chromatin loops and their anchors. This resolution is particularly important for detecting chromatin interactions that rely on transcription factors. To this end, we assessed chromatin contacts during atrial CM differentiation by Hi-C 3.0, an iteration of in situ Hi-C that is optimized for high-resolution detection of loops in addition to accurate detection of larger structures such as TADs and compartments (*26*). We collected two biological replicates from key cardiac differentiation time points, including pluripotency (d0), cardiac mesoderm (d2-d4), cardiac precursors (d6) and various CM stages (d11, d20 and d45) (Fig. 1A). At d45, ∼90% of cells expressed the CM marker cardiac muscle troponin T (cTnT+) as assessed by flow cytometry, indicating highly efficient CM differentiation. We sequenced libraries to an average depth of 1.69 billion raw read pairs and 761 million unique cis-interactions per time point, which allowed us to reach a resolution of 5 kilobases (kb) (fig. S1A). Biological replicates were highly reproducible and replicates were pooled for downstream analysis (fig. S1B,C). To assess correlations between chromatin contacts and gene expression, we also generated bulk RNA-sequencing libraries at each time point.

**Fig. 1.**
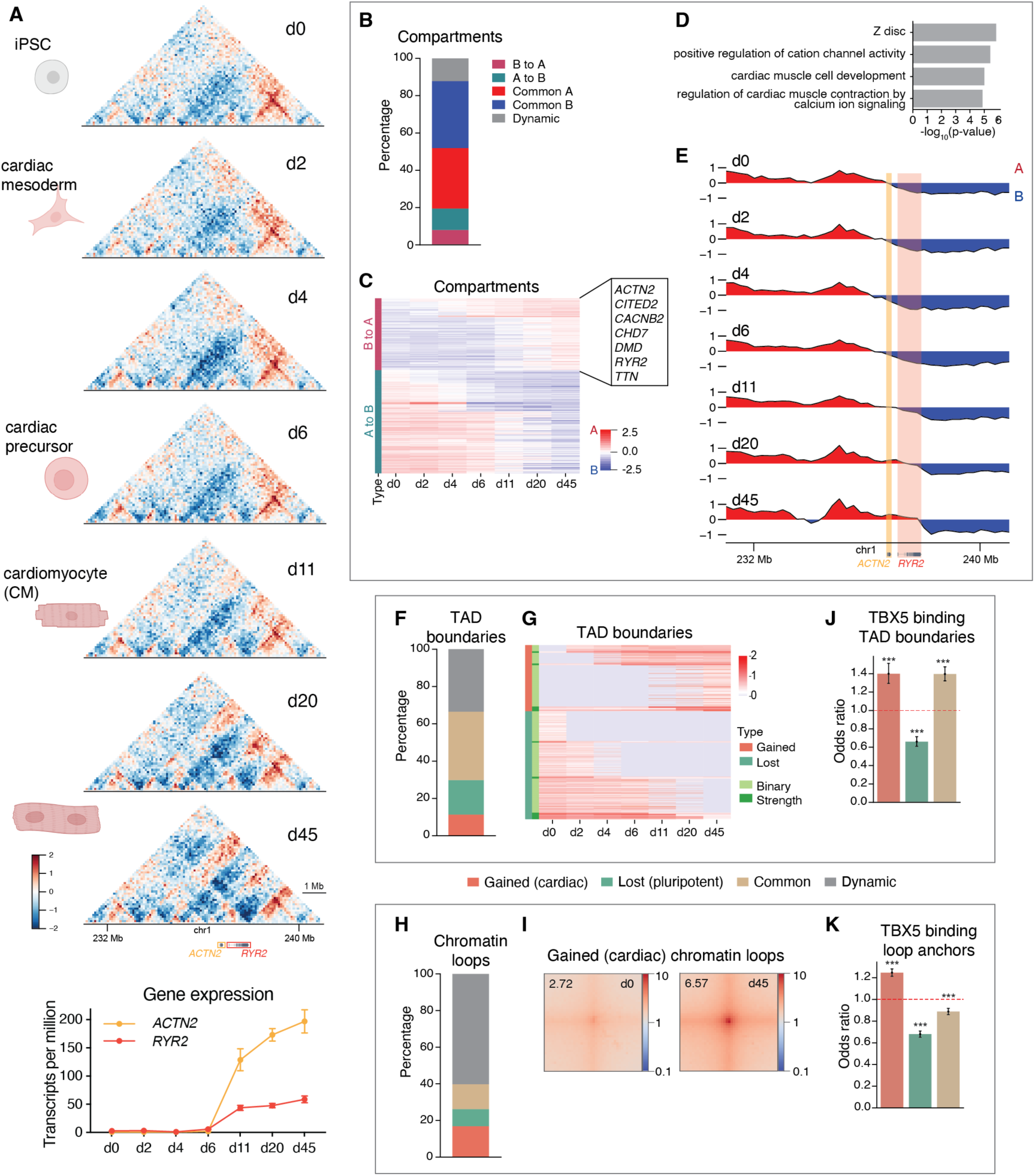
Dynamic reorganization of human cardiac chromatin. (**A**) Observed/expected contact maps for a 10 Mb region around *ACTN2* and *RYR2* during atrial cardiac differentiation, including RNA-seq gene expression (mean ± S.E.M.). Red and blue indicate regions with more or fewer contacts than expected, respectively. (**B**) Percentage of compartments that switch or remain the same during differentiation. (**C**) Heatmap of compartments switching from A to B or B to A. (**D**) Gene Ontology analysis of genes located in compartments switching from B to A after d11, highly correlated (Pearson R ≥ 0.8) with compartment scores. **(E**) Principal component scores showing progressive transition of *ACTN2* and *RYR2* loci from B to A compartments during differentiation. (**F**) Percentage of TAD boundaries that are common or change. (**G**) Heatmap of TAD boundaries that change during differentiation. (**H**) Percentage of chromatin loops that are common or change. (**I**) Aggregate Peak Analysis of loops gained in CMs (d45) v. iPSCs (d0). (**J,K**) Odds ratio plots of TBX5 binding at gained, lost and common TAD boundaries (J) or loop anchors (K). Two merged biological replicates per time point. Odds ratios compare each subset to all others; dotted lines at OR = 1 indicate no enrichment. ***p<0.001.

Consistent with other studies (*24, 25*), we observed large-scale chromatin reorganization during cardiac differentiation (Fig. 1A). We called A/B compartments for each time point and found that most of the genome (68.3%) remained in the same compartment throughout differentiation. Some compartments (19.5%) switched once from A to B or B to A and a smaller number (12.2%) switched multiple times (Fig. 1B,C). Focusing on genes with expression highly correlated with cardiac specific A compartments (d11 onwards, Pearson R ≥ 0.8), our GO analysis showed terms associated with heart morphogenesis and cardiac function (Fig. 1D). This included a number of essential cardiac genes, such as *TTN*, *DMD*, *RYR2*, *ACTN2*, *CITED2*, *CACNB2* and *CHD7*. Many of these are large genes encoding structural or contractile cardiac proteins that were insulated within B compartments in non-cardiac cells at earlier time points. We observed that A compartments expanded to cover these genes as cells differentiated (Fig. 1E). Increased gene expression sometimes preceded these chromatin changes (Fig. 1A,E; fig. S1D), suggesting that lineage specific transcriptional regulation and activity is a driver of compartment switching.

TADs were more dynamic than compartments, with only 36.6% of TADs shared across all stages of cardiac differentiation (Fig. 1F,G). Chromatin loops were even more dynamic, as only 13.5% were shared among all time points (Fig. 1H). This finding is consistent with observations in other cell types that suggest that chromatin loops form in a highly cell-type specific manner (*12*). We identified a total of 54,381 chromatin loops in CMs, a fifth of which (10,639) were specific to d11-d45 CMs compared to earlier time points (Fig. 1H; fig. S1E). As expected, CM specific chromatin loops had stronger interactions in CMs compared to pluripotent cells (Fig. 1I).

To understand whether TBX5 is involved in regulating cardiac-specific 3D chromatin, we assessed TBX5 binding during differentiation and correlated its binding profile with changes in 3D chromatin. For this, we engineered a biotin tagged-TBX5 WT iPSC line and assessed TBX5 binding by ChIP-seq. We found that TBX5 binding was enriched to similar degrees at both CM-specific TAD boundaries and TAD boundaries common to all time points (Fig. 1J). In contrast, TBX5 binding was more enriched at CM specific chromatin loop anchors than at loop anchors common to all time points (Fig. 1K). These results suggest that TBX5 may play a role in establishing chromatin loops in a CM-specific manner.

Together, these data highlight that the human genome is dynamically reorganized across all scales, from compartments, TADs, to chromatin loops, as pluripotent cells are directed towards the CM cell lineage.

### Cardiomyocyte chamber-specific subtypes display distinct 3D chromatin organization

Within the human heart, TBX5 is expressed in both atrial and ventricular CMs (*27*). Atrial and ventricular CMs have distinct functional and electrophysiological properties (*28, 29*) and can be distinguished transcriptionally (fig. S2A,B). It is possible that these differences are driven by distinct 3D genome organization. To explore this possibility, we generated a Hi-C 3.0 time course of ventricular CM differentiation. As with atrial differentiation, we observed dynamic changes in compartments, TADs and chromatin loops between stages of ventricular differentiation (fig. S3A-F). We specifically compared atrial d45 to ventricular d23 CMs, at which point cells had reached similar differentiation maturity in both protocols (fig. S2B), and found approximately 8.1% of compartments, 33.4% TADs and 48.1% chromatin loops were cell-type specific (fig. S3G-I). Thus, compared to the widespread dynamic changes in 3D chromatin from pluripotency to CMs during cardiac differentiation, there are more similarities in chromatin contacts between the two CM types. Nevertheless, we found that the expansion of A compartments across large structural protein-encoding genes such as *TTN* and *RYR2* began slightly earlier in the ventricular CM differentiations, indicating some differences in timing of identity acquisition at the chromatin level (fig. S3J-L). Supporting this hypothesis, ventricular CMs at d6 also appeared to be slightly more advanced transcriptionally than d6 atrial CMs (fig. S2B).

To determine if the two CM types could be distinguished by their 3D chromatin contacts, we differentiated iPSCs into either atrial or ventricular CMs and profiled chromatin contacts and DNA methylation in individual cells using single-cell methyl-Hi-C-seq (snm3C-seq) (*30, 31*). Single-cell resolution allowed us to assess the heterogeneity in 3D chromatin organization within and between CM differentiations. We generated snm3C-seq libraries for two biological replicates at three time points for each differentiation: iPSCs (d0), cardiac precursors (d6 for both atrial and ventricular differentiations) and late CMs (d23 for ventricular, d45 for atrial, which consisted of ∼89% and ∼81% cTnT+ cells, respectively).

After sequencing, we obtained on average 451,095 chromatin contacts per nucleus for 3,304 nuclei passing quality control. We clustered cells based on their chromatin contacts and observed that cells from atrial and ventricular differentiations formed distinct clusters, at both cardiac precursor and CM stages (Fig. 2A), thus confirming that atrial and ventricular CMs can be distinguished by genome-wide 3D chromatin organization. Clustering by DNA methylation or integrating DNA methylation with chromatin contacts revealed temporal differences (fig. S4A). However, all cardiac precursors (d6) clustered together and differences between atrial and ventricular populations only emerged at later stages of differentiation. This suggests that atrial versus ventricular CM fates are not driven by changes in methylation to the same extent as changes in 3D chromatin. We used Leiden clustering, a graph-based algorithm that optimizes the grouping of connected data points, and identified 11 distinct clusters that clearly separated the different time points and differentiation types from each other (Fig. 2B, fig. S4B,C). Focusing on the clusters with the largest number of cells, we could see changes in chromatin contacts around cardiac function genes, such as *ACTN2/RYR2* and *TTN,* between cardiac precursors and late CMs in both differentiations (Fig. 2C; fig. S4D). In addition, we identified differences in contacts around the atrial specific gene *KCNJ3* and the ventricular gene *IRX4* when comparing atrial to ventricular cells (Fig. 2D). Overall, these results show that atrial and ventricular CMs can be distinguished by their 3D genome early during differentiation and that these differences persist at more mature stages.

**Fig. 2.**
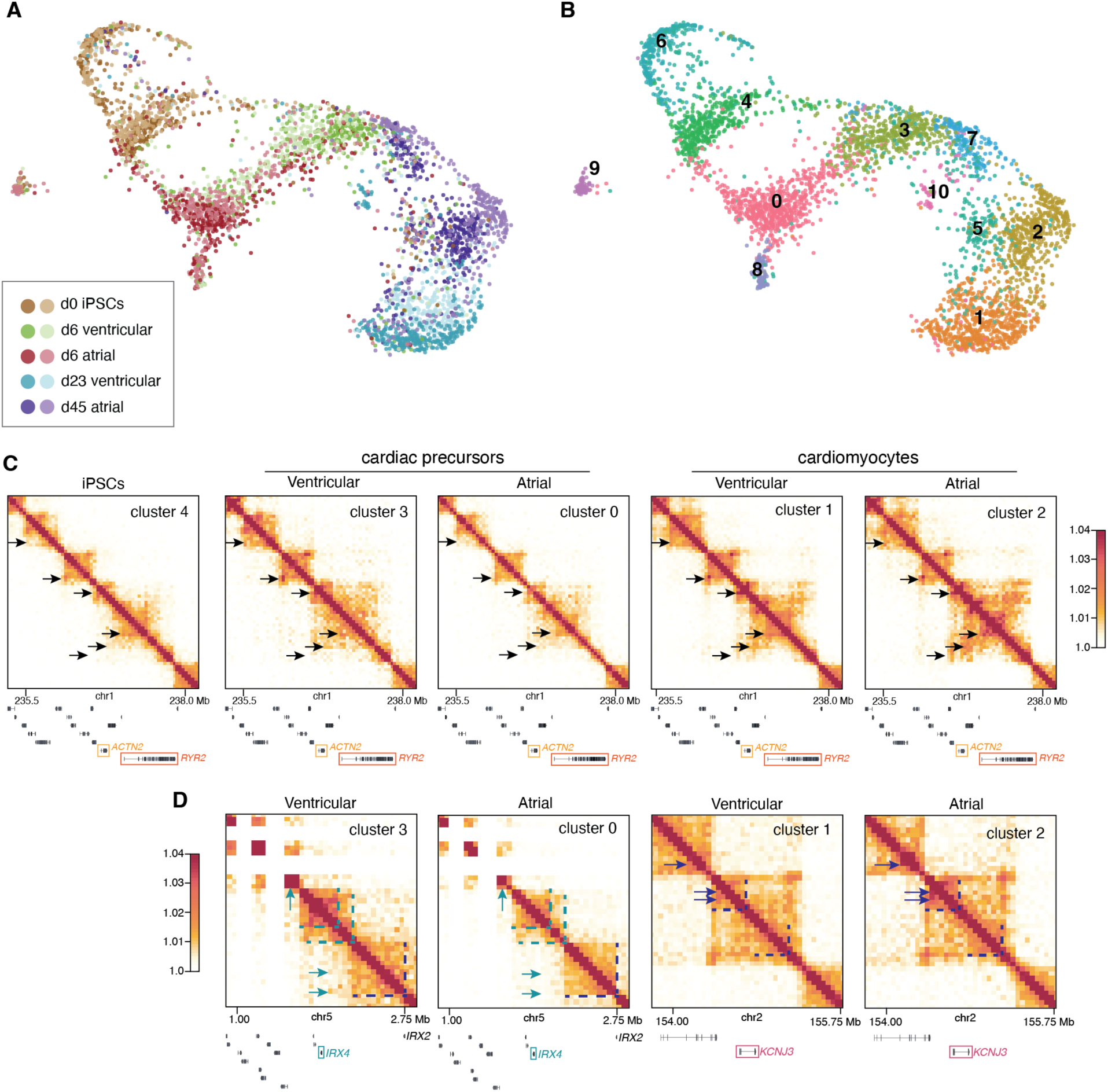
Cardiomyocyte subtypes display distinct 3D chromatin. (**A**) UMAP of atrial and ventricular differentiations, where the features are derived from Fast-Higashi embeddings of Hi-C contacts at 1 Mb resolution. Cells are colored by time point and cell type with the two shades indicating biological replicates. (**B**) Same UMAP as in (A), but with cells colored by Leiden clusters. (**C**) Imputed contact matrices of Leiden clusters 0-4 around cardiac enriched genes *ACTN2* and *RYR2*. Black arrows indicate regions of increased contact frequency in cardiomyocyte clusters (1 and 2). **(D**) Imputed contact matrices of ventricular (d23) and atrial (d45) cardiomyocyte enriched clusters around atrial-enriched gene *KCNJ3* and ventricular enriched-gene *IRX4*. Arrows and dashed lines indicate regions of increased contact frequency in atrial (navy) and ventricular (teal) cardiomyocytes. Scale in (C) and (D) shows log2(value+1), where value = balanced contact frequency. Two biological replicates per time point were analyzed and referred to in this figure.

### TBX5 influences chromatin organization at multiple scales in a dose-dependent manner

Since we found that TBX5 was enriched at cardiac specific chromatin loop anchors and to some extent TAD boundaries (Fig. 1J,K), we asked whether it may organize cardiac chromatin. For this, we performed Hi-C 3.0 in our previously generated cell lines (*20*), representing a *TBX5* allelic series (WT: *TBX5^+/+^*, heterozygous: *TBX5^in/+^*and null: *TBX5^in/del^*) (Fig. 3A). *TBX5^in/+^* CMs display sarcomere disarray indicating contractile dysfunction. Electrophysiological defects observed in human patients are recapitulated in *TBX5^in/+^*cells in the form of prolonged calcium transients. Both of these phenotypes are exacerbated in *TBX5^in/del^* CMs (*20*).

**Fig. 3.**
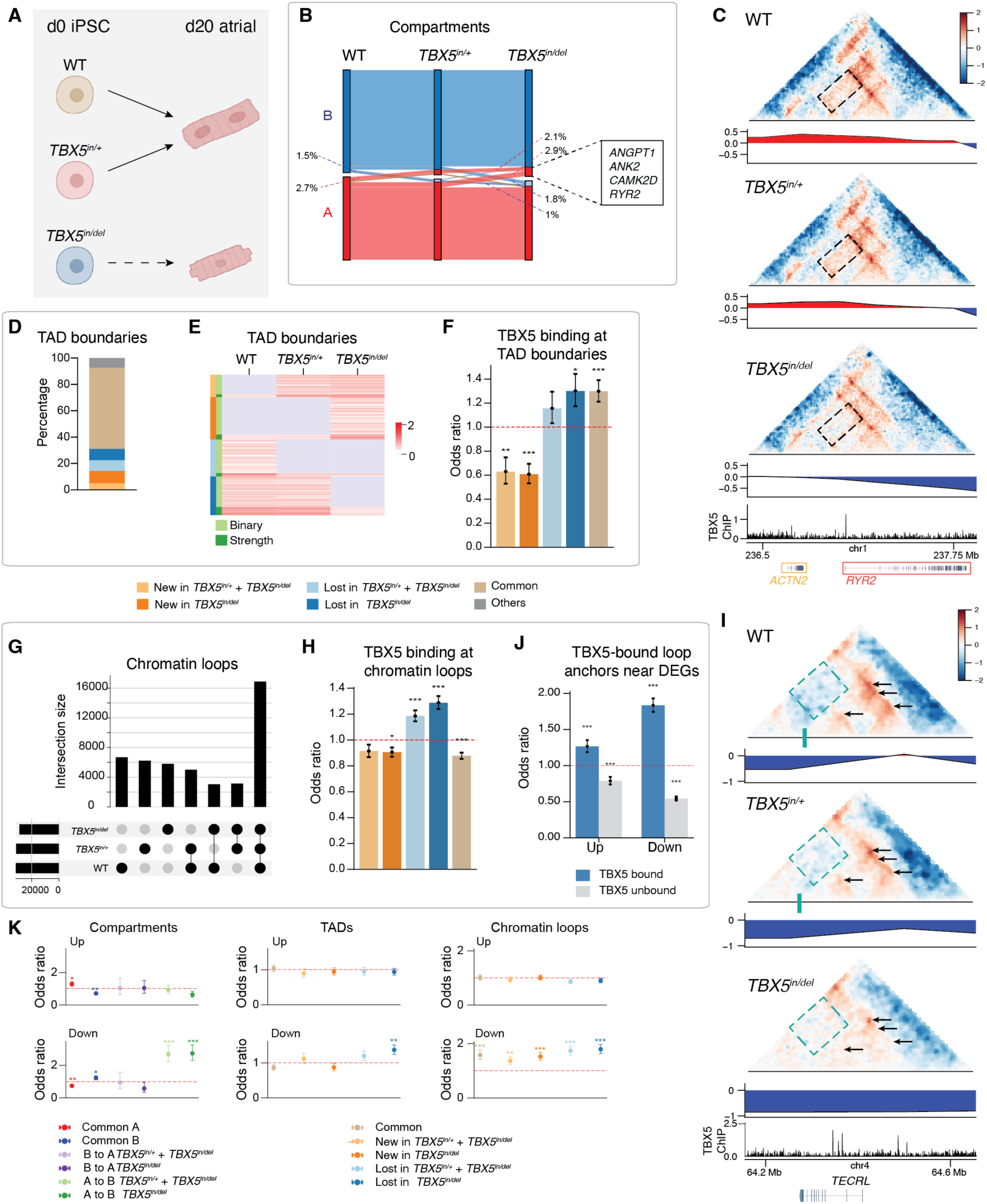
TBX5 dose-dependently influences 3D chromatin organization. (**A**) WT, *TBX5^in/+^* and *TBX5^in/del^* iPSC-derived atrial CMs. (**B**) Sankey plot showing compartment A/B switching. (**C**) Observed/expected contact map with A/B compartment principal components and TBX5 ChIP around *ACTN2* and *RYR2*. Boxed area indicates decreased contact frequency in *TBX5^in/del^* cells. (**D**) Percentage of TAD boundaries that are common or genotype-specific. (**E**) Heatmap of gained or lost TAD boundaries. **(F**) Odds ratio of TBX5 binding at TAD boundaries. (**G**) Upset plot of chromatin loops, indicating consensus loops between two or more genotypes. (**H**) Odds ratio of TBX5 binding at loop anchors. (**I**) Observed/expected contact map with A/B compartment principal components and TBX5 ChIP around *TECRL*. Green bar indicates the TAD boundary in WT and *TBX5^in/+^*. Boxed area indicates increased contact frequency in *TBX5^in/del^* cells. Arrows indicate lost loops in *TBX5^in/+^* and/or *TBX5^in/del^*cells. **(J**) Odds ratio of up- or downregulated genes near TBX5 bound loop anchors. (**K**) Odds ratio shows the association of up- or downregulated genes with common or changed compartments, TADs and chromatin loops. Two merged biological replicates per genotype. Odds ratios compare each subset to all others; dotted lines at OR = 1 indicate no enrichment. *P<0.05, **P<0.01, ***P<0.001.

We generated libraries for Hi-C 3.0 for two biological replicates of WT, *TBX5^in/+^* and *TBX5^in/del^* cells at d20 of atrial CM differentiation. As assessed by flow cytometry, the WT CMs were ∼76% cTnT+. The *TBX5^in/+^* line consistently produced smaller cTnT+ populations and the samples we collected for Hi-C 3.0 contained ∼62% cTnT+ cells. *TBX5^in/del^* iPSCs rarely differentiate into CMs and only ∼10% of the cells we collected were cTnT+ (fig. S5A). Despite the low number of CMs in *TBX5^in/del^* samples, they can occasionally be seen beating although they never have normal CM morphology. We sequenced Hi-C 3.0 libraries to an average depth of 1.29 billion raw read pairs and 706 million unique cis-interactions per genotype (fig. S5B), providing 5 kb resolution. The two *TBX5^in/+^* samples differed in their percentage of long cis contacts, which may reflect the phenotypic variation of heterozygous cells (fig. S5C). Nevertheless, *TBX5^in/+^* samples were overall more similar to each other than to WT and we pooled biological replicates for quantitative analysis (fig. S5D).

We observed clear, dose-dependent compartments changes in *TBX5^inl+^*and *TBX5^in/del^* cells. In *TBX5^in/+^* cells, compared to WT cells, a small fraction of compartments switched from A to B (2.7%) or from B to A (1.5%). In *TBX5^in/del^* cells, the frequency of these switches nearly doubled (4.9% and 2.8%, respectively) (Fig. 3B). The overlap in compartments that switched in both *TBX5^inl+^*and *TBX5^in/del^* cells highlighted regions that were particularly sensitive to TBX5 dosage (Fig. 3B). Overall, there was a small dose-dependent increase in the total number of B compartments in *TBX5^in/+^* and *TBX5^in/del^* CMs (fig. S6A). We clustered genomic regions based on compartment strength across the three genotypes and identified five clusters that were significantly different with clear dosage-sensitive patterns (fig. S6B). Compartments that switched from A to B in a TBX5 dose-dependent manner were associated with a significant enrichment for cardiac Gene Ontology terms and included important CM genes such as *ANGPT1, ANK2, CAMK2D and SCN10A* (Fig. 3B; fig. S6C). In addition, *ACTN2* and *RYR2* had switched back to B compartments in *TBX5^in/del^* cells, indicating that the expansion of cardiac A compartments depends on TBX5 (Fig. 3C). These findings highlight the importance of TBX5 for maintaining the correct spatial regulation of lineage-defining genes.

Reduced dosage of TBX5 also led to major changes in TAD organization and looping. We found that ∼38% of all TBX5 binding sites were within 10 kb of a TAD boundary or loop anchor (12% at TAD boundaries, 35% peaks at loop anchor). Among WT TADs, 17.3% were not detected in either *TBX5^in/+^*or *TBX5^in/del^* cells, and both *TBX5^in/+^* and *TBX5^in/del^* cells showed genotype-specific TADs. Overall, ∼40% of TADs identified across the genotypes were either lost or gained in *TBX5^in/+^*and/or *TBX5^in/del^* cells compared to WT (Fig. 3D,E). In total, 10.05% of all WT TAD boundaries were bound by TBX5 in d20 atrial CMs. TAD boundaries that were lost in *TBX5^in/+^* and/or *TBX5^in/del^* cells were highly likely to be bound by TBX5 in WT cells (Fig. 3F). TAD boundaries that were common to all three genotypes were also enriched for TBX5 binding, suggesting that only some TAD boundaries depend on TBX5 binding for their formation or maintenance. In contrast, the boundaries of the gained TADs in *TBX5^in/+^* and/or *TBX5^in/del^*cells tended not to be bound by TBX5 in WT cells (Fig. 3F). Therefore, we defined TADs or chromatin loops as “TBX5 sensitive” if they were lost in *TBX5^in/+^*and/or *TBX5^in/del^* samples. To complement our TBX5 ChIP dataset, we generated CTCF ChIP-seq data in each of the three genotypes. Although many TAD boundaries were lost, only ∼11% of WT CTCF binding sites were lost in the *TBX5^in/+^* and/or *TBX5^in/del^* cells and most of them did not overlap with WT TAD boundaries (fig. S6D). Since many TAD boundaries were sensitive, mechanisms other than loss of CTCF binding likely regulate these TAD changes. Indeed, this supports our hypothesis that many CM TAD boundaries depend on TBX5. Surprisingly, the main impact of reduced TBX5 dosage on CTCF binding was gain, rather than loss, of binding sites, with ∼23% of all analyzed CTCF binding sites being novel in *TBX5^in/del^* cells (fig. S6E).

Since TBX5 binding was enriched at CM specific chromatin loops, we predicted that the major impact of reducing TBX5 would be on chromatin loops. In total, 15.43% of WT loop anchors were bound by TBX5 in d20 atrial CMs. Indeed, of all the loops identified across all three genotypes, up to 50% were specific to *TBX5^in/+^*and *TBX5^in/del^* cells (Fig. 3G). As with gained TADs, the anchors of these gained loops were not enriched for TBX5 binding in WT cells (Fig. 3H), indicating they are regulated by TBX5-independent or downstream mechanisms. By contrast, TBX5 binding was enriched at the anchors of loops lost in *TBX5^in/+^* and *TBX5^in/del^* cells (Fig. 3H), indicating that chromatin loops are especially vulnerable to reduced dosage of TBX5. GATA4, another cardiac TF known to occupy loci and co-regulate transcription with TBX5 (*32–35*), was also enriched at TBX5-sensitive chromatin loops (fig. S6F). Many chromatin loops, including at the *TBX5* locus, spanned TAD boundaries (fig. S6G). A number of chromatin loops with an anchor at the *TBX5* promoter or gene body were bound by TBX5, either at the anchor or within the looped region, in WT CMs. Some of these loops were lost in *TBX5^in/+^* CMs, and all of them were lost in the *TBX5^in/del^*CMs, suggesting that TBX5 regulates its own 3D genome architecture and possibly gene expression (fig. S6H).

Dose-dependent changes in chromatin organization at all scales are exemplified around the TBX5 target gene *TECRL*, whose expression is known to depend on TBX5 dosage (*20*). In the WT CMs, the *TECRL* gene body is within a weak B compartment, and a ∼50 kb weak A compartment lies upstream of the promoter region. The strength of the B compartment gradually increased as the TBX5 dosage decreased, whereas the weak A compartment was completely abrogated by the loss of only one copy of *TBX5* (Fig. 3I). The TAD boundary that insulates the *TECRL* TAD in both WT and *TBX5^in/+^* CM was lost in *TBX5^in/del^*cells, leading to increased contact frequency between neighboring TADs (boxed area in figure). Many chromatin loops were identified with anchors inside the *TECRL* gene body as well as between the promoter and the upstream euchromatic region in WT CMs. The majority of these chromatin loops were lost in *TBX5^in/+^* CMs and all were lost in *TBX5^in/del^* cells. We found TBX5 binding sites were enriched within chromatin loops anchored at the *TECRL* promoter, suggesting that the presence of TBX5 is required at these sites for correct 3D genome organization (Fig. 3I).

### Reduced transcription of TBX5 target genes is linked to altered 3D chromatin

Given the transcriptional dysregulation associated with reduced TBX5 levels (*20*) (fig. S7), we also wanted to understand whether transcriptional changes were linked to changes in 3D chromatin organization and TBX5 binding. Genes that were downregulated by a reduction in TBX5 dosage were highly enriched near chromatin loop anchors with TBX5 binding (Fig. 3J). Interestingly, both gained and lost loops were associated with downregulated genes, suggesting that changes to looping dynamics in general are associated with loss of transcription (Fig. 3K). Downregulated genes were also highly enriched within compartments that underwent an A to B switch, and in TADs that were lost in *TBX5^in/+^* and/or *TBX5^in/del^*cells (Fig. 3K). These findings are consistent with TBX5 acting as a direct activator of the transcription of these genes.

In contrast, upregulated genes in *TBX5^in/+^* and/or *TBX5^in/del^* cells were not enriched at chromatin structures that changed (Fig. 3K). TBX5-bound loop anchors were enriched near upregulated genes, but not to the same extent as downregulated genes (Fig. 3J). This suggests that TBX5 binding tends to mediate the activation of target genes rather than their repression. TBX5 has been shown to act as a transcriptional repressor by recruiting the NuRD complex at a number of genes (*36*). It is possible that some upregulated genes are repressed by TBX5 but others may be due to secondary effects such as the downregulation of other genes or a redistribution of TFs that normally cooperate with TBX5, leading to ectopic gene expression, as previously shown (*35*).

### Heterogeneity of *TBX5* dosage-sensitive 3D chromatin

Variations in chromatin architecture due to loss of TBX5 may not affect all cells uniformly. In particular, the differences in the percentage of long cis contacts observed in our two *TBX5^in/+^*samples highlight possible heterogeneity that is masked in bulk samples. To look for any genotype specific heterogeneity, we also generated snm3C-seq data for WT, *TBX5^in/+^* and *TBX5^in/del^* cells at d20 of atrial CM differentiation (n = 2). Samples were ∼74% cTnT+ in the WT, ∼62% cTnT+ in *TBX5^in/+^* and ∼18% cTnT+ in *TBX5^in/del^*samples (fig. S8A). We obtained on average ∼1.5 million reads and 761,900 chromatin contacts per nuclei for 2,170 nucleus passing quality control.

We clustered cells using chromatin contacts and/or by methylation, identifying clear differences between WT and *TBX5^in/+^*cells compared to *TBX5^in/del^* cells (Fig. 4A; fig. S8B,C). Similar to our atrial and ventricular time course data, we found the separation by methylation was less distinct than for chromatin contacts (fig. S8B). Focusing on chromatin contact clusters, we performed Leiden clustering and found WT and *TBX5^in/+^* samples were somewhat similar (mostly falling in clusters 0 and 2), whereas *TBX5^in/del^*samples clustered separately (clusters 1, 3 and 5). (Fig. 4A,B; fig. S8D). Cluster 4, which contained a mix of the three genotypes, is likely a non-CM population as it showed distinctly less contact frequency around the cardiac genes *ACTN2* and *RYR2* (fig. S8E). At the TBX5-dosage sensitive gene *TECRL*, we found that the two clusters containing a majority of WT and *TBX5^in/+^*cells showed increased contacts around the gene compared to the clusters containing mostly *TBX5^in/del^* cells (Fig. 4C). There were also differences in contact frequency between the two WT and *TBX5^in/+^* enriched clusters (green arrows in Fig. 4C). At *ANK2* and *CAMK2D,* other genes that showed dose-dependent changes in chromatin contacts in *TBX5^in/+^* and *TBX5^in/del^* cells (Fig. 3B), we saw changes in contacts between WT/*TBX5^in/+^* clusters and *TBX5^in/del^* clusters (Fig. 4D). Again, there were also some differences between clusters of the same genotype, showing that single cells display heterogeneity in 3D organization. Overall, these contact maps show that TBX5 dosage sensitivity explains some of the differences in 3D chromatin organization in single cells.

**Fig. 4.**
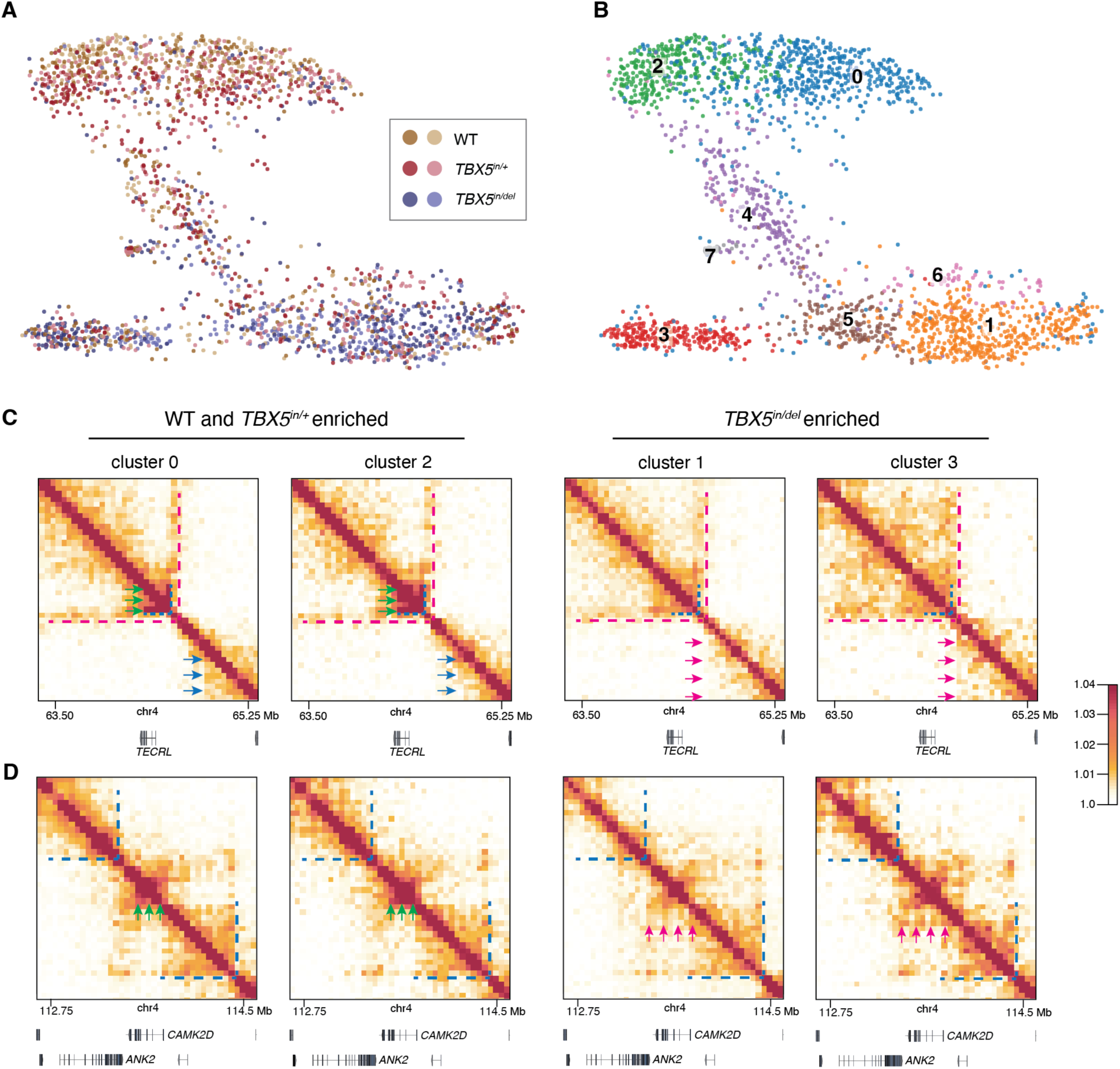
Chromatin contact heterogeneity associated with reduced dosage of *TBX5*. (**A**). UMAP of cells from the TBX5 allelic series, where the features are derived from Fast-Higashi embeddings of Hi-C contacts at 1 Mb resolution. Cells are colored by TBX5 genotype, with the two shades indicating biological replicates. (**B**) Same UMAP as in (A), but with cells colored by Leiden clusters. (**C,D**) Imputed contact matrices of WT and *TBX5^in/+^* enriched clusters (cluster 0, 2) and *TBX5^in/del^* enriched clusters (cluster 1, 3) around TBX5-dosage sensitive genes *TECRL, ANK2* and *CAMK2D*. Cluster specific annotations colored by clusters in (B), where cluster 3 is recolored as magenta. Green and blue arrows indicate differences between the two WT and *TBX5^in/+^*enriched clusters, magenta arrows indicate differences between the two *TBX5^in/del^*enriched clusters. Blue and magenta dashed lines indicate differences between WT/*TBX5^in/+^* enriched clusters and *TBX5^in/del^* enriched clusters. Scale shows log2(value+1), where value = balanced contact frequency. Two biological replicates per genotype were analyzed and referred to in this figure.

### TBX5 and CTCF co-occupy cardiac specific TADs and chromatin loops

Since TBX5 binding was enriched at both TAD boundaries that were lost in *TBX5^in/+^* and *TBX5^in/del^* cells as well as at TAD boundaries shared across all genotypes (Fig. 3F), this suggested that only some TBX5-bound TAD boundaries are sensitive to loss of TBX5.

To explore the reasons behind this differential sensitivity, we assessed co-occupancy by TBX5 and CTCF (fig. S9A-E). Genome wide, ∼26% of TBX5 binding sites overlapped with CTCF binding (fig. S9F). Almost all (95%) of TBX5-bound TAD boundaries were co-occupied by CTCF (fig. S9A,C). Although TAD boundaries bound by TBX5 alone were a small proportion of all TBX5-bound boundaries, most of them were sensitive to loss of TBX5 (fig. S9A), indicating that TBX5 is directly required to maintain these TADs. In contrast, CTCF and TBX5 co-occupied boundaries were underrepresented as sensitive to TBX5 dosage reduction compared to TBX5-only boundaries (18%, relative risk 0.26, p<0.0001) (fig. S9A), suggesting that the presence of CTCF may protect TADs from loss of TBX5. In general, we found that TBX5-sensitive TADs had lower insulation scores compared to insensitive TADs. This was particularly evident for TBX5-only bound TAD boundaries, which showed lower insulation scores compared to CTCF and TBX5 co-bound ones, including at insensitive TAD boundaries (Fig. 5A). Many critical cardiac transcriptional activators including *HAND2, MYOCD, NR2F1, NR2F2 and TBX3* were inside or nearby CTCF and TBX5 co-bound sensitive TADs (Fig. 5B; fig. S9A).

**Fig. 5.**
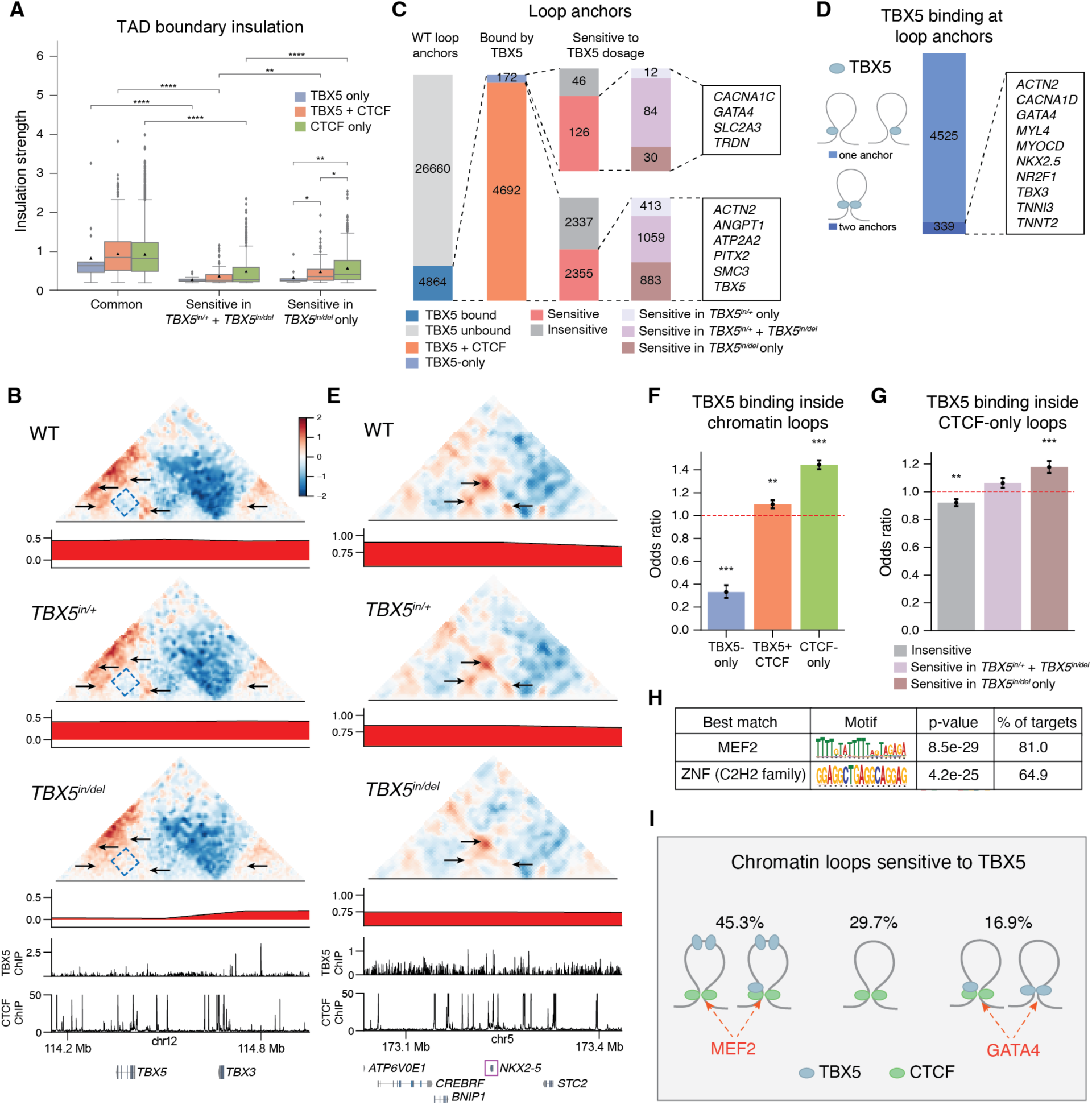
TBX5 establishes cardiac chromatin contacts through both CTCF-dependent and independent mechanisms. (**A**) Insulation strength of common and sensitive TAD boundaries separated by TBX5/CTCF binding status. (**B**) Observed/expected contact maps, principal component scores highlighting A/B compartments, WT TBX5 and CTCF peaks around *TBX5* and *TBX3.* Arrows indicate reduced contact frequency in *TBX5^in/+^* and/or *TBX5^in/del^* cells. Boxed area indicates increased contact frequency in *TBX5^in/del^* cells. (**C**) Percentage of TBX5-bound loop anchors segregated into TBX5-only or CTCF+TBX5 co-occupied loops, and by sensitivity to loss of TBX5. Gene list indicates those near sensitive contacts. (**D**) Proportion of all chromatin loop anchors bound by TBX5 separated by those that are bound at one or both anchors. **(E**) Same legend as (B), around *NKX2-5.* Arrows indicate reduced contact frequency in *TBX5^in/+^* and/or *TBX5^in/del^* cells. (**F**) Odds ratio of TBX5 binding within loop anchors bound by TBX5 or CTCF. (**G**) Odds ratio of TBX5 binding within sensitive CTCF-only bound loop anchors. **(H**) Motif analysis of loop anchors for CTCF-only bound sensitive loops. (**I**) Summary of TBX5-sensitive chromatin loops. Two merged biological replicates per genotype. Odds ratios compare each subset to all others; dotted lines at OR = 1 indicate no enrichment. *p<0.05, **p-value<0.01, ***p<0.001, ****p<0.0001.

Similar to TAD boundaries, loop anchors bound by TBX5 alone were rare but more sensitive to loss of TBX5 compared to TBX5 and CTCF co-bound anchors (relative risk 1.46, p<0.0001) (Fig. 5C). Only ∼7% of all TBX5-bound loop anchors had TBX5 at both anchors (Fig. 5D). Interestingly, these loops were enriched for genes encoding other cardiac lineage-determining TFs *GATA4, NKX2-5* and co-activator *MYOCD*, in addition to key heart contraction genes *TNNT2, TNNI3, ACTN2* and *MYL4 (*Fig. 5D,E). Given that these loops are particularly sensitive to TBX5 dosage, failure to normally regulate these genes likely explains why *TBX5^in/+^* and *TBX5^in/del^*cells display dose-dependent decreases in CM differentiation efficiency. Thus, TAD boundaries and chromatin loops bound by TBX5 binding alone or co-bound by CTCF and TBX5 but with weak insulation are sensitized to loss of TBX5.

### TBX5 and CTCF co-binding tends to occur in A compartments leaving B compartments more sensitive to loss of TBX5

Since there was a relatively large number of compartment changes (∼8% of genome in *TBX5^in/del^* cells) (Fig. 3B), we wanted to know whether changes in TBX5 binding at TAD boundaries or loop anchors related to larger scale compartment changes. TBX5 binding sites were more enriched within A compartments (fig. S10A), consistent with the notion that TBX5 is largely a transcriptional activator. Similarly, TBX5 and CTCF co-binding were enriched in A compartments, but depleted in B compartments at both TAD boundaries and loop anchors (fig. S10B). TBX5-sensitive TADs were not enriched within compartments that switched in *TBX5^in/+^* and *TBX5^in/del^*cells (fig. S10C), suggesting that TBX5-sensitivity at TAD boundaries is not the driver of compartment switching. Interestingly, TBX5-sensitive TADs were predominantly within B compartments common to all genotypes (fig. S10D), which may be sensitive as a result of less CTCF co-occupancy than in A compartments. In contrast to TADs, TBX5-sensitive loops were most enriched in compartments that switched from A to B (fig. S10C,D). We found that GATA4 tended to be bound together with TBX5 in both A and B compartments (fig. S10E). Therefore, although large-scale compartment switching is not completely explained by changes to smaller scale structures like TAD boundaries and some chromatin loops, these results further support the notion that CTCF co-binding with TBX5 generally buffers regions from reduced TBX5 dosage.

### Mechanisms that sensitize CTCF-only bound loops to TBX5

While TBX5 binding was enriched at TAD boundaries and loop anchors, the majority of TADs and chromatin loops that were sensitive to TBX5 loss did not have TBX5 binding sites at their boundaries or anchors (fig. S10F). We called these CTCF-only loops.

TBX5 is enriched at active enhancer regions (*33, 35*). Active enhancers are emerging as regulators of cohesin recruitment that enable cohesin-dependent loop extrusion and loop formation (*37, 38*) and are known to be functionally influenced by TFs (*39*). We hypothesized that CTCF-only bound loop anchors may be impacted in two ways in the absence of TBX5: firstly, by reduced cohesin loading at TBX5-bound enhancers leading to less insulation of these loops, and secondly, by failed recruitment of other factors that might be important in allowing CTCF to function at loop anchors.

To address the first hypothesis, we asked whether TBX5 binding was enriched within sensitive loops, where TBX5 may regulate cohesin complex loading at some distance from the loop anchor. Overall, ∼57% of cardiac loops contained TBX5 binding sites. At TBX5-sensitive loops with CTCF-only anchors, we found that TBX5 binding within loops was enriched compared to other sensitive and insensitive loops (Fig. 5F,G), especially compared to TBX5-only bound loop anchors. These data support the requirement for TBX5 binding within sensitive loops to enable loop formation, possibly by facilitating cohesin loading.

To address the second hypothesis, we analyzed motifs at non-TBX5 bound sensitive loop anchors and found enrichment for MEF2 and ZNF (C2H2 family) motifs (Fig. 5H). Of the cardiac expressed MEF2 family proteins, both *MEF2A* and *MEF2C* were expressed by a large proportion of CMs and dysregulated in the absence of TBX5, with *MEF2A* showing dose-dependent downregulation in *TBX5^in/+^* and *TBX5^in/del^* cells (fig. S7). This analysis suggests that 3D chromatin organization may also be sensitive to MEF2A loss downstream of TBX5. Additionally, *Tbx5* and *Mef2c* genetically interact in mouse heart development (*20*), and are part of a set of TFs that can reprogram fibroblasts to CMs (*40*). Therefore, MEF2A or MEF2C are likely additional cardiac TFs with a putative role in regulating 3D chromatin.

In summary, we find that TBX5 affects 3D chromatin organization through both cis and trans mechanisms. In cis, TBX5 binding is directly required to form numerous cardiac-specific chromatin loops and TADs with and without CTCF co-occupancy. In trans, TBX5 may have multiple roles that impact 3D chromatin organization, including regulating the chromatin architecture around and expression of other TFs that may have additional, and as yet unknown, roles in 3D chromatin regulation (Fig. 5I).

## Discussion

We identified the disease-linked, lineage-specific cardiac TF TBX5 as a direct dose-dependent regulator of human CM 3D organization. TBX5 is enriched at TAD boundaries and chromatin loops that emerge during CM differentiation. We discovered that diverse features of the 3D genome—chromatin loops, TAD boundaries and compartments—are sensitive to reduced TBX5 dosage. Of relevance to human disease, *TBX5* heterozygous cardiomyocytes had considerable alterations in 3D chromatin folding, which was further exacerbated by the complete loss of *TBX5*. This indicates that chromatin organization is highly sensitive to TBX5 dosage. We observed two major TBX5-dependent mechanisms. In most regions sensitive to the loss of TBX5, TBX5 is bound within chromatin loops. These loops are regulated by CTCF at anchors enriched for motifs for MEF2 TFs, which interact genetically with TBX5 (*20*). In a smaller number of sensitive regions, TBX5 binds directly to the loop anchor or TAD boundary. We found that co-occupancy of CTCF at TBX5-bound loop anchors and TAD boundaries reduces the sensitivity of the loops or TADs to reduced TBX5 dosage, with many of these structures remaining unchanged even in the complete absence of TBX5. By contrast, co-occupancy with GATA4, another cardiac TF, does not appear to protect 3D chromatin from loss of TBX5, as GATA4 predominantly associated with and co-occupied TBX5 binding sites at sensitive regions. TBX5 and GATA4 interdependently bind and co-regulate cardiac gene expression programs (*33, 35*), and their association together at regions required for cardiac specific 3D chromatin looping suggests an additional aspect of their joint function in gene regulation. Finally, a subset of regions were sensitive to the loss of TBX5 through indirect mechanisms. Our results point to a complexity of TF-TF interactions in the regulation of 3D chromatin during development.

We propose that TBX5 regulates cardiac 3D chromatin organization through multiple, distinct but functionally overlapping mechanisms. First, TBX5 binding at loop anchors suggests that TBX5 is directly required to anchor cardiac-specific loops. Multiple mechanisms have been proposed for TF mediated chromatin looping, including direct or cofactor oligomerization, interactions with structural factors and recruitment of transcription and remodeling machinery (*41*). Since the majority of anchors are co-occupied by CTCF, we propose that TBX5 may act as a cell-type determinant of where loops should be created, with CTCF then structurally insulating them. TBX5’s role in this may occur through recruitment of chromatin remodeling complexes (*42, 43*), histone modifying enzymes (*44*) or other co-factors. Since many TBX5 binding sites overlap with CTCF binding sites, it is also possible that the presence of TBX5 is directly required for CTCF binding at those sites in CMs. At the few loops where TBX5 is bound at both anchors, TBX5 may have a direct role in loop formation, possibly through homodimer formation to bring together distant loci (*45*). Second and more frequently, cardiac chromatin loops are associated with TBX5 binding sites within looped regions. It is possible cell-type specific cohesin loading at enhancers is regulated by lineage-specific TFs (*39*) and that in CMs TBX5 may play that role. Indeed, modeling predicts that a distinct set of TFs are required for chromatin folding in different cell types (*46*). Finally, there may be effectors downstream of TBX5 that regulate 3D genome organization. Loss of TBX5 leads to the dysregulation of multiple cardiac TFs, which may have as-yet-unknown roles in regulating chromatin contacts.

Many but not all changes in 3D chromatin caused by loss of TBX5 were associated with dysregulated gene expression. TAD boundary strength correlates with active transcription (*47*), and the TBX5-sensitive TAD boundaries were generally weak, suggesting they are not associated with active transcription. Together, these observations support the hypothesis that TBX5 regulates 3D chromatin independently of transcription, via structural and cohesin-regulating roles. Given the dynamic reorganization of the 3D genome we observed during CM differentiation, it is possible that the transcriptional consequences of lost contacts at some regions would be evident by assessing earlier stages in differentiation as shown in datasets for other non-cardiac lineage-restricted TFs MYOD and IKAROS (*13, 17*). Since our allelic series datasets capture a single time point it is unknown at which stage during differentiation these 3D contacts are established, although clearly TBX5 is important for maintaining them.

In disease contexts, known pathogenic alterations in 3D chromatin contacts impact one or a few gene loci, particularly through structural variants that impact the expression of a salient disease-linked gene (*48*). Within the context of the heart, this research is currently limited. Recently, a large genetic deletion in patients that disrupts a TAD boundary and *PITX2* regulation was associated with electrophysiology defects (*49*). Our results highlight large scale, disease-relevant mechanisms through which TBX5 haploinsufficiency may lead to dysregulated gene expression in patients with CHDs.

Mechanisms of haploinsufficiency are poorly understood. One possible mechanism involves changes to chromatin accessibility (*50*). Here we have described an additional layer: alterations in 3D genome organization. Given the preponderance of haploinsufficient TFs as regulators of developmental processes dysregulated in human syndromes (*2*), these extensive chromatin changes might reflect a more generalizable mechanism of developmental defects caused by TF haploinsufficiency.

## Materials and Methods

### Maintenance of iPS cells and differentiation to cardiomyocytes

Protocols for use of iPSCs were approved by the Human Gamete, Embryo and Stem Cell Research Committee, as well as the Institutional Review Board at UCSF. WTc11 iPSCs (gift from Bruce Conklin, available at NIGMS Human Genetic Cell Repository/Coriell #GM25256) were used in this study. WTc11 iPSCs with genome-edited mutations for *TBX5* (*TBX5^in/+^* and *TBX5^in/del^*) were previously generated and characterized in both ventricular (*20*) and atrial cardiomyocyte (CM) differentiations (Bruneau lab unpublished). WTc11 iPSCs used for atrial and ventricular time course Hi-C 3.0 experiments and corresponding RNA-seq were grown on hESC-qualified Matrigel (Corning #354277). Human iPSCs for all other experiments were grown on growth factor-reduced basement membrane Matrigel (Corning #356231) in mTeSR-1 (StemCell Technologies #85850) or mTeSR+ medium (StemCell Technologies #100-0276). iPSCs were routinely passaged using ReLesR (StemCell Technologies #100-0483). For CM differentiations, iPSCs were dissociated using Accutase (StemCell Technologies #07920) and seeded into 6-well or 12-well plates. Cells were grown for three days until 70-90% confluency and induced using Stemdiff Cardiomyocyte Ventricular (StemCell Technologies #05010) or Atrial (StemCell Technologies #100-0215) kits, according to manufacturer’s instructions.

### Flow cytometry

During CM harvest for Hi-C 3.0, scMethyl-HiC or ChIP, about 5% of each sample was kept for flow cytometry analysis of CM differentiation efficiency, as assessed by proportion of cardiac troponin T2+ cells. Cell pellets were resuspended in 4% formaldehyde (from 16% stock concentration, methanol-free, ThermoFisher Scientific #28906) and incubated on nutator for 15 min at room temperature. Samples were centrifuged for 5 min at 200 x g at 4°C. Samples were washed twice in DPSB and stored at 4°C for up to 1 week before preparing for flow cytometry. Cells were permeabilized in FACS buffer (0.5% w/v saponin, 4% FBS in PBS) for 15 min at room temperature. Cells were stained with a mouse monoclonal antibody against cardiac isoform Ab-1 Troponin (ThermoFisher Scientific #MS-295-P, final concentration 5 µg/ml) or the IgG1kappa isotype control (ThermoFisher Scientific #14-4714-82) for 1 hour at room temperature, gently vortexing cells every 20 min. Samples were washed with FACS buffer and the stained with donkey anti-rabbit IgG Alexa 488 (ThermoFisher Scientific #A21206, 1:200 dilution, final concentration 5 µg/ml) for 1 hour at room temperature, gently vortexing cells every 20 min. Cells were washed with FACS buffer and filtered into tubes with 35-micron mesh caps (Corning #352235). 10 mg/ml DAPI (Invitrogen #D1306) was added to final concentration of 1 µg/ml. At least 10 000 cells were sample were analyzed using the BD LSRFortessa X-20 (BD Bioscience) and final results were processed using FlowJo (FlowJo, LLC).

### Hi-C 3.0 library preparation and sequencing

Cells were grown in 3 wells of a 6-well dish per biological replicate, with all 3 wells pooled together after dissociation for the crosslinking step (approximately 10 million cells). iPSCs were dissociated in Accutase (StemCell Technologies #07920) for 5 min. Differentiated cells were dissociated in 0.25% Trypsin-EDTA (5 min for d2, d4 or d6 cells; 10 min for d11, d20, d23, d45 CMs; Gibco #25200056), followed by quenching in 15% FBS in DPBS. Cells were lifted off the dish and resuspended to single cells by repeatedly using P1000 pipette, then transferred to a 15 ml tube and centrifuged at 800 rpm for 5 min at room temperature. Pellets were washed once in 5 ml HBSS before resuspending entire cell pellet in exactly 10 ml HBSS. 16% methanol-free formaldehyde (ThermoFisher Scientific #28908) was added to a final concentration of 1% and samples incubated on nutator at room temperature for 10 min. Samples were quenched with 2.5 M glycine to a final concentration of 0.125 M and incubated on nutator at room temperature for 5 min, then 15 min at 4°C and followed by centrifugation at 1000 x g for 5 min at room temperature. Samples were washed once in DPBS. Pellets were resuspended in 1 ml of 3 mM DSG in PBS, transferred to 1.5 ml tubes and incubated on nutator for 40 min. Samples were quenched with 2.5 M glycine to a final concentration of 0.125 M and incubated on nutator at room temperature for 5 min. Samples were centrifuged at 2000 x g for 5 min at RT. Cell pellets were resuspended in 5 mg/ml BSA in DPBS, to prevent cells sticking to tube after DSG crosslinking, and centrifuged at 2000 x g for 5 min at 4°C. Supernatant was carefully removed and cell pellets were snap frozen in liquid nitrogen and stored at -80°C.

Two biological replicates per time point were processed for Hi-C 3.0 and library preparation (with the exception of ventricular d2 and d4 where n = 1 was used). Samples were processed for Hi-C using Arima-HiC+ kit (Arima Genomics). >1 µg per sample was used as determined by Arima’s Estimating Input Amount Protocol using Qubit Fluorometer and dsDNA HS Assay Kit (ThermoFisher Scientific #Q32854).

Samples were processed as per Arima’s user guide for mammalian cell lines with one exception: DNA was digested using 50 U DpnII (New England Biolabs #R0543M) and 50 U DdeI (New England Biolabs #R0175L) restriction enzymes in 1x NEB Buffer 3.1 for 60 min at 37°C. All samples passed Arima-QC1 Quality Control. Sequencing libraries were prepared as per Arima’s user guide for Library Prep Using KAPA Hyper Prep Kit (Roche #7962347001). Between 1.5-4 µg of DNA was fragmented in 130 µl microtube (Covaris #520045) using a Covaris S2 sonicator for 55s at 10% duty cycle, intensity 4, 200 cycles per burst in a water bath maintained at 4°C. An average fragment size of 400 bp was verified using Agilent 2100 Bioanalyzer. Between 400-1000 ng of size-selected DNA was used for biotin enrichment, end repair, dA-tailing and adapter ligation. The number of PCR cycles was determined following Arima-QC2 Quality Control using KAPA Library Quantification Kit (Roche KK4824) and libraries were amplified to at least 10 nM. All samples passed Arima-QC2 Quality Control. DNA cleanup steps were done using Ampure XP beads (Beckman Coulter #A63881). Final libraries were quantified using Qubit Fluorometer and dsDNA HS Assay Kit and assessed on Agilent BioAnalyzer. Pooled, equimolar libraries were sequenced using NovaSeq technology (Illumina). All atrial time course samples were sequenced in one batch, all TBX5 allelic series were in one batch and ventricular time course samples were sequenced in two batches (one biological replicate per batch) using 150 bp paired-end reads.

### Hi-C 3.0 data processing

Hi-C 3.0 data were processed using 4D Nucleome (4DN) Hi-C data Processing Pipeline (v0.2.7) (https://data.4dnucleome.org/resources/data-analysis/hi_c-processing-pipeline), which generated .mcool file with iterative correction and eigenvector decomposition (ICE) normalization and .hic file with Knight-Ruiz (KR) normalization. For the analyses that do not explicitly compare replicates, data from biological replicates for each timepoint or genotype were merged together to generate a combined Hi-C matrix for high resolution after checking their reproducibility using HiCRep (*51*). Compartments and TAD boundaries were called using cooltools from .mcool files (*52*) at the resolution of 250 kb and 5 kb, respectively. HiCCUPS (*53*) and Peakachu (*54*) were used to detect chromatin loops at both 5 kb and 10 kb resolution from .mcool files and .hic files, respectively. The loops called by each tool at the two resolutions were first merged and then the resulting loops were further combined to get a union set of loops by removing the ones with both anchors within 20 kb of another one.

### A/B compartment analyses

Compartments for each group of samples (atrial or ventricular time course, or *TBX5* allelic series) were classified as common A, common B, B to A, A to B and dynamic. Compartments that consistently remained in the A or B state across all time points or genotypes were classified as common A or common B, respectively. For the time series data, compartments that switched from B to A at a specific time point and remained in the A compartment for all subsequent time points were classified as B to A (“cardiac specific”). Conversely, those that switched from A to B at a given time point and remained in B were classified as A to B. For the atrial time course data, to evaluate whether the compartment change is ahead of gene expression change or the other way, the earliest time point for compartment change was identified by finding the time point at which the compartment switched to a different status and kept the status for all subsequent time points.

For the *TBX5* allelic series, compartments that were in the B compartment in WT samples and shifted to A in *TBX5^in/+^* and *TBX5^in/del^*samples, or those in B in both WT and *TBX5^in/+^* but switched to A in *TBX5^in/del^*, were classified as B to A. Similarly, compartments that changed from A to B across genotypes were classified as A to B. All other compartments exhibiting changes across time points or genotypes, but not fitting these categories, were classified as dynamic.

Differential compartment analyses for the *TBX5* allelic series was performed using dcHiC (*55*), which identified compartments that had statistically significant changes in their compartment scores, regardless of changes between the A and B states. For each group of samples (atrial/ventricular time course, *TBX5* allelic series), we used the compartments called as previously described. The compartment score was determined by the first eigenvector from Principal Component Analysis (PCA). Next, dcHiC quantile normalized the compartment scores across samples. Then, a distance score (based on Mahalanobis distance) is calculated for each compartment per genotype based on these normalized compartment scores. This distance score represents the degree of compartment score change for one compartment compared to all compartments within the *TBX5* allelic series. An FDR threshold of 0.01 was used to determine significant differential compartments. K-means was used to cluster these significant compartments.

### TAD analyses

The .bigwig files generated by cooltools during TAD boundary calling were converted to .wig format using bigWigToWig. Regions lacking insulation scores were identified and extracted as gap regions for each sample. For each group of samples, these TAD boundaries and gap regions were combined by merging those within 10 kb of each other using bedtools merge, respectively. The presence of each TAD boundary in a sample was identified using bedtools intersect. TAD boundaries were filtered out from further analyses if the TAD regions formed by a boundary and its adjacent boundaries overlapped with gap regions in more than 60% of the samples containing that boundary.

For the time course data, the TAD boundaries were categorized into four groups: common, gained, lost and dynamic. TAD boundaries that either appeared (binary) or showed higher insulation strength (log2FC > 0.6) at a specific time point and remained consistent in all subsequent time points were classified as gained (“cardiac specific”). Conversely, boundaries that disappeared (binary) or had lower insulation strength (log2FC < -0.6) from a certain time point onwards were classified as lost. TAD boundaries that were present across all time points but did not meet the criteria for gained or lost were categorized as common. All other boundaries that exhibited changes but did not fit into these categories were classified as dynamic. For the *TBX5* allelic series data, TAD boundaries were similarly classified as (1) common, (2) gained in *TBX5^in/+^* and *TBX5^in/del^*, (3) gained in *TBX5^in/del^*, (4) lost in *TBX5^in/+^* and *TBX5^in/del^*, (5) lost in *TBX5^in/del^*, and (6) dynamic, based on their binary presence or changes in insulation strength (log2FC > 0.6 or log2FC < -0.6).

The relative risk analysis of TBX5 and CTCF co-bound sensitive TADs was calculated by the percentage of TBX5+CTCF co-bound TAD boundaries that are lost in *TBX5^in/+^*and/or *TBX5^in/del^* (17.83%) divided by the percentage of TBX5-only bound TAD boundaries that are lost in *TBX5^in/+^* and/or *TBX5^in/del^* (68.63%).

### Chromatin loop analyses

Similar to the TAD boundaries, a union set of chromatin loops for each group was generated by removing the ones with both anchors within 20 kb of another one. The presence of a chromatin loop in each sample was identified using pgltools intersect. For the time series data, the chromatin loops were categorized as common, gained (“cardiac specific”), lost and dynamic others. For the *TBX5* allelic series, the chromatin loops for the TBX5 allelic series were classified as (1) common, (2) gained in *TBX5^in/+^* and *TBX5^in/del^*, (3) gained in *TBX5^in/del^*, (4) lost in *TBX5^in/+^* and *TBX5^in/del^*, (5) lost in *TBX5^in/del^*, and (6) dynamic others, based on their binary presence.

Aggregate peak analysis (APA) for the cardiac-specific loops were performed using coolpup.py and visualized using plotpup.py. The relative risk analysis of TBX5 and CTCF co-bound sensitive loops was calculated by the percentage of TBX5-only bound loop anchors that are lost in *TBX5^in/+^* and/or *TBX5^in/del^* (73.26%) divided the percentage of TBX5+CTCF co-bound loop anchors that are lost in *TBX5^in/+^* and/or *TBX5^in/del^*(50.19%).

### snm3C-seq library preparation and sequencing

Cells were grown in one well of 6-well dish per biological replicate. iPSCs were dissociated in Accutase (StemCell Technologies #07920) for 5 min. Differentiated cells were dissociated in 0.25% Trypsin-EDTA (5 min for d2, d4 or d6 cells; 10 min for d11, d20, d23, d45 CMs; Gibco #25200056), followed by quenching in 15% FBS in DPBS. Cells were lifted off the dish and resuspended to single cells by repeatedly using P1000 pipette and then transferred to a 15 ml tube and centrifuged at 300 x g for 5 min at room temperature. Pellets were washed once in 1 ml DPBS and counted using Countess II Automated Cell Counter with optimized setting. 2 million cells were aliquoted into a 1.5 ml tube and centrifuged at 300 x g for 5 min at room temperature. Cell pellets were resuspended in 1 ml DBPS and 16% methanol-free formaldehyde (ThermoFisher Scientific #28908) was added to final concentration of 2% and incubated on nutator at room temperature for 5 min. Samples were quenched with 2.5 M glycine to a final concentration of 0.2 M and incubated on nutator at room temperature for 5 min, followed by centrifugation at 1000 x g for 5 min at 4°C. Samples were washed once in ice cold DPBS and cell pellets were snap frozen in liquid nitrogen and stored at -80°C.

Two replicates per time point were processed for snm3C-seq and library preparation. Cells were subjected to in situ 3C procedure following the steps in the Arima-3C BETA Kit (Arima Genomics), with some modifications. Cells were lysed for 15 min on ice in 20 µl of Lysis Buffer. 24 µl of Conditioning Solution was added, and cells were transferred to a 0.2 mL PCR tube, and mixed gently by pipetting. Tubes were incubated at 62°C for 30 min in a thermal cycler, with the lid temperature set to 85°C. 20 µl of Stop Solution 2 was added, and mixed gently by pipetting. Tubes were incubated at 37°C for 15 min in a thermal cycler with a lid temperature set to 85°C. 28 µl of the restriction enzyme digestion mix (9.8 µl water, 9.2 µl Buffer H, 4.5 µl Enzyme H1, 4.5 µl Enzyme H2) was added and gently pipette mixed. Tubes were incubated in a thermal cycler for 37°C for 60 min, 65°C for 20 min, then 25°C for 10 min. 82 µl of ligation mix was added (70 µl Buffer C, 12 µl Enzyme C) and gently mixed by pipetting, then incubated at 25°C for 15 min. Nuclei were transferred to a tube containing 850 µl 1% BSA in DPBS on ice and filtered through a 30um Celltrics strainer to a new tube on ice. Nuclei were centrifuged at 1000 x g for 10 min at 4°C (in bucket rotor). Supernatant was removed and pellets were resuspended each sample in 800 µl of 1% BSA in DPBS. Nuclei were stained with DRQA7 at 1:200 and mixed gently with a pipette and incubated on ice for 5 min. One tube of Proteinase K (Zymo #D3001-2-D) was dissolved in 1 mL of Proteinase K Resuspension Buffer (Zymo #D3001-2-B) then added to a mix of 15 mL M-Digestion Buffer 2x (Zymo #D5021-9) and 14 mL water to make pK Digestion Buffer. 2 µl of pK Digestion Buffer were added to each well of an Eppendorf twin.tec® PCR Plate 384 LoBind®, skirted, 45 µl, PCR clean (Eppendorf #0030129547). Single DRQA7 positive nuclei were sorted into individual wells of the 384-well plate using a Sony SH800 cell sorter. One 384-well plate was used per biological replicate. Plates were sealed then incubated in a thermal cycler at 50°C for 20 min with a heated lid at 85°C for nuclei digestion.

### Library preparation and Illumina sequencing for snm3C-seq

The snm3C-seq bisulfite conversion procedure and library preparation protocol was carried out as previously described (*56, 57*). This procedure was automated using the Beckman Coulter Biomek i7 liquid handlers for all reactions in 384 and 96-well plates. The snm3C-seq libraries were shallow sequenced on the Illumina NovaSeq 2000 to 2-10 M reads to check quality. Successful libraries were deeply sequenced on the NovaSeq6000 or NovaSeqX Plus instruments using 150 bp paired-end reads.

### Mapping and analysis of snm3C-seq

Raw sequence FASTQ files were mapped using YAP (Yet Another Pipeline) software (cemba-data v1.6.9), as previously described (*57*). Briefly, FASTQ files were demultiplexed by cell barcodes and read quality assessment (cutadapt, v.2.10). Two-pass mapping was performed with bismark (v0.20, with bowtie2 v2.3) for alignment to hg38 reference genome assembly. BAM file processing and QC was performed using samtools (v1.9) and Picard (v3.0.0). Chromatin contacts were called and methylome profiles were generated using Allcools software (v1.0.23).

Single-cell methylation analyses were performed with the *ALLCools* suite (*56*). Cells were first filtered based on their mapping rates (>50) and methylation fraction at mCG (>0.5), mCH (<0.2) and mCCC (<0.03) sites. For the atrial and ventricular time course datasets, 3304 nuclei (from total 3840 nuclei) passed quality control filters. For *TBX5* allelic series datasets, 2170 nuclei (from total 2304 nuclei) passed quality control filters. To cluster single-cell, the fraction of mCH and mCG methylation in 100 kb bins across the genome were used. These bins were filtered by coverage (>50 and <3000), and sites in ENCODE blacklist regions were not considered. The top 20000 most variable regions with a support vector regression (SVR) were selected. Next, Principal Component Analysis was performed on these top regions for dimensionality reduction. The top PCs were visualized with UMAP, and cells were colored according to their time point, cell type, and/or genotype.

Single-cell Hi-C analyses were performed with *Fast-Higashi* (*58*). In brief, the sparse single-cell Hi-C maps per cell were imputed with a partial random walk with restart algorithm for 1 Mb bins across each chromosome. Next, the imputed scHi-C tensors are decomposed to produce cell embeddings and chromosome-specific bin weights, transformation, and meta-interaction matrices. 256-dimension embeddings were obtained to represent each cell and used for downstream visualization and clustering. Leiden clustering was performed for cells using the single-cell methylation and single-cell Hi-C PCs and embeddings. The Leiden clusters were then used to pseudo bulk the scHiC contacts. With *cooler* (*59*), the contacts from cells in each cluster were merged at 50kb resolution, and the resulting bulk matrices were balanced with ICE normalization. Finally, to integrate the single-cell methylation and single-cell HiC clusters, the respective principal components and embeddings were concatenated.

### Bulk RNA-seq library preparation

For the atrial differentiation, samples from WT cells were collected at d0 (iPSCs), d2, d4, d6, d11, d20 and d45 for three biological replicates. For the ventricular differentiation, samples from WT cells were collected at d6, d10, d12, d15, d30 for three biological replicates. Cells were grown in 1 well of a 6- or 12-well dish per biological replicate. iPSCs were dissociated in Accutase (StemCell Technologies #07920) for 5 min. Differentiated cells were dissociated in 0.25% Trypsin-EDTA (5 min for d2, d4 or d6 cells; 10 min for d11, d20, d23, d45 CMs; Gibco #25200056), followed by quenching in 15% FBS in DPBS. Cells were lifted off the dish and resuspended to single cells by repeatedly using P1000 pipette and then transferred to a 1.5 ml tube and centrifuged at 300 x g for 5 min at room temperature. Pellets were snap frozen in dry ice. Total RNA was isolated using Qiagen RNeasy micro kit (Qiagen #74004) with QIAshredder (Qiagen #79656) and eluted in 25 µl water. RNA concentration was determined using Nanodrop and quality was checked using Agilent Bioanalyzer. Atrial CM RNA-seq libraries were generated using Illumina Stranded Total RNA Prep with Ribo-Zero Plus kit were prepared as per manufacturer’s instructions using 150 ng of input RNA. Ventricular CM RNA-seq libraries were generated using NuGEN Ovation Ultralow System V2 kit as per manufacturer’s instructions. Library concentration was quantified using Qubit Fluorometer and dsDNA HS Assay Kit and quality assessed on Agilent Bioanalyzer. Libraries were pooled and sequenced on a NovaSeq X (Illumina) using 50 bp paired-end reads for atrial differentiation samples or on a NextSeq500 using 75 single-end reads for ventricular differentiation samples.

### Bulk RNA-seq analysis

The .fastq files were analyzed using nf-core Nextflow RNA-seq pipeline (DOI: 10.5281/zenodo.1400710) (*60*) aligning to hg38 reference genome and using STAR for alignment (*61*) and Salmon quantification (*62*). Differential gene expression analyses were performed for each pair of timepoints using DESeq2 (*63*). The time point that a gene shows increased (log2FC≥1, adjusted p-value<0.05) or decreased (log2FC≤-1, adjusted p-value<0.05) expression compared to previous time points and remained consistent in all subsequent time points was considered as the earliest point for gene expression changes.

### Single cell RNA-sequencing

Single cell RNA-seq was performed in the *TBX5* allelic series at d20 atrial CM differentiation for two biological replicates and available from GSE285169 From differential gene expression, genes that showed increased or decreased expression in both *TBX5^in/+^* and *TBX5^in/del^* or *TBX5^in/del^* alone were identified to investigate the association of gene expression with 3D chromatin changes.

### Generation of TBX5-bio cell line

A codon-optimized BioTAP (*64*) tag was inserted immediately after the start methionine codon in the N-terminus of the endogenous TBX5 locus using ribonucleoprotein (RNP) complexes of sgRNA (GCCCTCGTCTGCGTCGGCCA, ordered from Synthego) and SpCas9-NLS ‘HiFi’ protein (QB3 MacroLab, University of California Berkeley). This tag can be biotinylated by endogenous biotin ligases. The BioTAP tag was codon optimized for integration using HDR Alt-R gene blocks donor template (IDT). RNPs were prepared by mixing 1 µl of 40 µM Cas9 with 1.2 µl of 100 µM sgRNA and made to a total volume of 10 µl with Lonza P3 transfection buffer (Lonza #197179) and incubated at room temperature for 10 min.

250000 WTc11 iPSCs were resuspended in 12.2 µl Lonza P3 transfection buffer and nucleofected with 1 µg codon-optimized BioTAP donor template and 10 µl RNPs using Lonza Nucleofector 4D system in 96-well format using DS-138 settings. After nucleofection, mTeSR1 supplemented with CloneR (StemCell Technologies #05888) was added to cells and incubated for 10 min at 37°C, before transferring cells into a single well of a matrigel-coated 48-well plate. The following day, media was replaced with fresh mTeSR1. The cells were expanded into a 12-well plate. Single cells were dissociated with Accutase (StemCell Technologies #07920) and were sorted into 96-well plates for colony selection using Namocell Hana single-cell dispenser. Plates were cultured in mTeSR1 supplemented with CloneR for 3 days post sorting to ensure survival and then replaced with mTeSR1. Colonies were screened by PCR using primers pairs flanking the whole insertion site (F: 5’-TCCTCAGAGCAGAACCTTGC, R: 5’-CTTACCTGCTGGGTGAAGG), 5’ integration site (F: 5’-AAACTGCTCCCTCCTGTCAC, R: 5’-ATTTTCATGGCCTCAAGCAC) and 3’ integration site (F: 5’-CCGATGGTAAGGTCGAGAAG, R: 5’-CGAGGTCTCCTTACCTGCTG).

Amplicon from whole insertion site PCR was amplified and verified by Sanger sequencing. Copy number variation was determined by droplet digital PCR (Bio-Rad QX200 Droplet Reader) and using the RPP30 assay (Bio-Rad #10031243) and comparing to custom primers (F: 5’-GCGTTTATCTCCGTCTCCATTT and R: 5’-GGACTGTCAGTAAGATCCTTGTT) and probe (5’-CAAGCACCGTTTGCCCTGCTTTAA) for the knock-in TBX5-biotin allele. The cells were determined to have a normal karyotype.

### ChIP-seq library preparation

For TBX5 ChIP, WT TBX5-bio CMs were collected at d11 (n = 2), d20 (n = 6) and d45 (n = 4) during atrial differentiation. For CTCF ChIP-seq, WT, *TBX5^in/+^* and *TBX5^in/del^* CMs were collected at d20 of atrial differentiation (n = 2 per genotype). CMs were grown in one 6-well plate per replicate, with all 6 wells pooled together after dissociation for the crosslinking step (approximately 20 million cells). Each plate was one biological replicate. CMs were dissociated in 0.25% Trypsin-EDTA (Gibco #25200056) for 10 min, followed by quenching in 15% FBS in DPBS. CMs were lifted off the dish and resuspended to single cells by repeatedly using P1000 pipette and then transferred to a 15 ml tube and centrifuged at 800 rpm for 5 min at room temperature. Pellets were washed once in 10 ml DPBS plus 1x protease inhibitors (DBPS+PI) before resuspending entire cell pellet in exactly 10 ml DPBS+PI. 16% methanol-free formaldehyde (ThermoFisher Scientific #28908) was added to a final concentration of 1% and samples incubated on nutator at room temperature for 10 min. Samples were quenched with 2.5 M glycine to a final concentration of 0.125 M and incubated on nutator at room temperature for 5 min, followed by centrifugation at 500 x g for 5 min at 4°C. Samples were washed once in ice cold DPBS+PI, before being transferred to 2 ml protein lo-bind rubes, centrifuged at 2500 x g at 4°C. for 5 min and cell pellets were snap frozen in liquid nitrogen and stored at -80°C.

TBX5-bio cross-linked cells were thawed on ice and resuspended in 1 ml of ice cold cell lysis buffer (25 mM Tris-HCl, pH 7.4, 85 mM KCl, 0.25% Triton X-100 and 1x Protease inhibitors in ddH2O) and incubated at 4°C rotating for 30 minutes. Sample were centrifuged at 2500 x g for 5 min at 4°C and resuspended in 1 ml nuclear lysis buffer (0.5% SDS, 10 mM EDTA, 50 mM Tris-HCl, pH 8.0, and 1x protease inhibitors in ddH2O). Samples were transferred to 1 ml millitube (Covaris #520130) and sheared using a Covaris S2 sonicator for 15 min at 5% duty cycle, intensity 5, 200 cycles per burst in a water bath maintained at 4°C. After shearing, 5 µl of each sample was kept for reverse crosslinking and the remaining sonicated sample was transferred to a new protein lo-bind tube and snap frozen in liquid nitrogen. The 5 µl sample was made up to 100 µl and incubated for 30 min at 37°C with 1 µl RNase A, followed addition of 12 µl 5 M NaCl and reverse crosslinking with 1 µl proteinase K at 65°C overnight. The following day the reverse crosslinked samples were purified using Qiagen MinElute PCR Purification kit (Qiagen #28006) and DNA quantified using Qubit Fluorometer and dsDNA HS Assay Kit (ThermoFisher Scientific #Q32854). Samples were run on an Agilent 2100 Bioanalyzer to verify that chromatin was sheared to fragments between 100-500bp in size. Based on these measurements, the amount of DNA in each lysed sample could be extrapolated and equal amounts of sheared chromatin could be used per ChIP. Sheared samples were thawed quickly at 37°C and the equivalent of 55 µg of chromatin was transferred to a 5 ml protein lo-bind tube for ChIP and made up to 1 ml in nuclear lysis buffer. 50 µl (5%) of each sample was collected as input and stored at 4°C until reverse crosslinking step. ChIP samples were then diluted 1:5 with ChIP dilution buffer (16.7 mM Tris-HCl pH 8.0, 1.2 mM EDTA, 0.01% SDS, 1.1% Triton X-100, 167 mM NaCl and 1x protease inhibitors in ddH2O). 50 µl of pre-washed Dynabeads MyOne Streptavidin T1 beads (ThermoFisher Scientific #65601) were added to each sample and incubated overnight with rotation at 4°C. The following day, beads were collected using a magnetic stand and transferred to 2 ml protein lo-bind tubes and washed with 1 ml twice of the following wash buffers: wash buffer 1 (2% SDS in ddH2O), wash buffer 2 (0.1% sodium deoxycholate, 1% Triton X-100, 1 mM EDTA, 50 mM HEPES pH 7.5, 500 mM NaCl in ddH2O), wash buffer 3 (250 mM LiCl, 0.5% IGEPAL CA-630 [Sigma #I8896], 0.5% sodium deoxycholate, 1 mM EDTA and 10 mM Tris-HCl pH 8.0 in ddH2O) and TE buffer (10 mM Tris-HCl pH 7.5 and 1 mM EDTA ddH2O). After adding each wash buffer, beads were vortexed for 15 s, spun briefly and collected against a magnet. DNA was eluted with 150 µl elution buffer (1% SDS, 10 mM EDTA, 50 mM Tris-HCl pH 8.0 in ddH2O) shaking at 1000 rpm at 65°C overnight. The following day, beads were collected on a magnetic stand and supernatant was transferred to a new DNA lo-bind tube. Both ChIP and input samples were incubated at 37°C for 30 min at 37°C with 1.5 µl RNase A, followed addition of 18 µl 5 M NaCl and reverse crosslinking with 1.5 µl proteinase K at 65°C for 6 hours. DNA was purified using Qiagen MinElute PCR Purification kit (Qiagen #28006).

CTCF ChIP-seq was performed generally as above with changes detailed here. The following buffers were used: cell lysis buffer recipe (20 mM Tris-HCl pH 8.0, 85 mM KCl, 0.5% IGEPAL CA-630 and 1x protease inhibitors in ddH2O), nuclear lysis buffer recipe (1% SDS, 10 mM EDTA, 50 mM Tris-HCl, pH 8.0, and 1x protease inhibitors in ddH2O), wash buffer 1 recipe (20 mM Tris-HCl pH 8.0, 150 mM NaCl, 2 mM EDTA, 0.1% SDS and 1% Triton X-100 in ddH2O), wash buffer 2 recipe (20 mM Tris-HCl pH 8.0, 500 mM NaCl, 2 mM EDTA, 0.1% SDS and 1% Triton X-100 in ddH2O) and wash buffer 3 (250 mM LiCl, 1% IGEPAL CA-630 [Sigma #I8896], 1% sodium deoxycholate, 1 mM EDTA and 10 mM Tris-HCl pH 8.0 in ddH2O) and elution buffer (1% SDS, 100 mM NaHCO3 in ddH2O). 70 µg of chromatin was used per sample. After diluting in dilution buffer, each ChIP sample was incubated rotating at 4°C overnight with 6 µg antibody (combination of both Active Motif #61311 [discontinued] and Abcam #ab128873 used in each sample) and the following day incubated with 50 µl prewashed Magna ChIP protein A/G beads (Sigma #16-663) for 4 hours at 4°C. Bead washes were performed by resuspending beads in 1 ml buffer using P1000 pipette, spinning briefly and collecting on magnet.

Following the bead washes, DNA was eluted in 50 µl shaking at 37°C, transferring to new tube and repeating bead elution to pool eluates to a total volume of 100 µl. Both ChIP and input samples were incubated at 37°C for 30 min at 37°C with 1 µl RNase A, followed by addition of 12 µl 5 M NaCl and reverse crosslinking with 1 µl proteinase K at 65°C overnight. DNA was purified using ChIP DNA Clean and Concentrator kit (Zymo Research #D5205).

For Illumina sequencing library preparation, the entire ChIP sample or 2 µl of each input was used. End repair was performed in a 100 µl reaction with 15 U T4 DNA polymerase (New England Biolabs #M0203L), 5 U Klenow fragment DNA polymerase (New England Biolabs, #M0210L), 50 U T4 PNK (New England Biolabs #M0201L), 400 µM dNTP (Promega #U1511) and 1x T4 DNA ligase buffer w 10 mM ATP (New England Biolabs #B0202S) in nuclease-free water for 30 min at room temperature, followed by 1.6x bead:sample AmpureXP purification. Entire eluate was used for A-tailing in a 50 µl reaction with 1 mM dATP (New England Biolabs #N0440S), 15 U Klenow 3’>5’ exonuclease (New England Biolabs #M0212L) and 1x NEB buffer 2 in nuclease-free water for 30 min at 37°C, followed by 1.6x bead:sample AMPureXP purification. Entire eluate was used for adapter ligation in a 50 µl reaction with 6000 U T4 DNA ligase (New England Biolabs #M0202L), 20 nM annealed and uniquely indexed adapters and 1x T4 DNA ligase buffer with 10 mM ATP (New England Biolabs #B0202S) in nuclease-free water for 2 hours at room temperature, followed by 1x bead:sample AMPureXP purification. Adapters were prepared by annealing following HPLC purified oligos: 5’-AATGATACGGCGACCACCGAGATCTACACTCTTTCCCTACACGACGCTCTTCCGATC*T and 5’Phos-GATCGGAAGAGCACACGTCTGAACTCCAGTCACNNNNNNATCTCGTATGCCGTCTTCTGCTTG where * represents a phosphothiorate bond and NNNNNN is a Truseq index sequence.The entire eluate was used for PCR amplification in a 50 µl reaction with 1x NEB Next High-Fidelity 2x PCR master mix (New England Biolabs #M0541L), 10 µM primers (F: 5’-AATGATACGGCGACCACCGAGATCTACACTCTTTCCCTACACGA, R: 5’-CAAGCAGAAGACGGCATACGAGAT) in nuclease free water, using thermocycler settings 98°C for 30 sec; 16 cycles of 98°C for 10 sec, 58°C for 40 sec and 72°C for 30 sec; 72°C for 5 min and followed by 0.8x bead:sample AMPureXP purification. DNA was quantified using Qubit Fluorometer and dsDNA HS Assay Kit and quality assessed on Agilent Bioanalyzer. TBX5-bio ChIP-seq libraries were sequenced on a NovaSeq X (Illumina) using 50 bp paired-end reads. CTCF ChIP-seq libraries were sequenced on a NextSeq2000 (Illumina) using 50 bp paired-end reads.

### ChIP-seq analysis

Fastq files were aligned to the hg38 reference genome using bowtie2 (*65*). Reads were filtered to include those with a mapq score of 30 or greater and removing duplicate reads using SAMtools (*66*). Blacklisted regions were removed using BEDtools (*67*).

Chip-seq peaks were called on each individual replicate relative to input using macs2 and the narrowpeaks parameter (*68*). CTCF ChIP bigwig files for visualization were generated using deepTools2 bamCoverage (*69*).

To define CM TBX5 binding sites, the union set of peaks detected in d11, d20 and d45 samples was used. TBX5 ChIP bigwigs for visualization were generated as log2 fold change over input using deepTools2 bigwigCompare (*69*).

### GATA4 ChIP-seq analysis

GATA4 peaks from two biological replicates of human iPSC-CMs were downloaded from the GEO database (accession number: GSE85628). The coordinates were converted to hg38 from hg19 using liftover and the peaks from biological replicates were merged using bedtools merge. The occupancy of GATA4, TBX5, and CTCF at TAD boundaries and loop anchors were identified by intersecting TAD boundaries and loop anchors with their peaks, respectively.

### Motif enrichment analysis

To explore other factors that could regulate the loops sensitive to TBX5 loss, we applied MEME with classical mode (*70*) to identify the motifs enriched in anchors of CTCF-only bound loops that were lost in *TBX5^in/+^* and/or *TBX5^in/del^*. The discovered motifs were further compared with the known vertebrate TF PWMs in the JASPAR database using Tomtom (*70, 71*).

## Acknowledgments

We thank members of the Gladstone Stem Cell Core, Flow Cytometry Core and the UCSF Center for Advanced Technologies for their expert assistance. We thank Mylinh Bernardi of the Gladstone Genomics Core for assistance with bulk RNA-seq library preparation, Tatyana Sukonnik for generating ventricular CMs for RNA-seq libraries, Reuben Thomas of the Gladstone Bioinformatics Core for valuable feedback on bioinformatics analysis, Françoise Chanut for editorial assistance, and members of the Bruneau and Pollard labs for discussions and comments. The Gladstone Flow Cytometry Core is funded by NIH S10 RR028962, the James B. Pendleton Charitable Trust and DARPA. Sequencing was performed at the UCSF CAT, supported by UCSF PBBR, RRP IMIA, and NIH 1S10OD028511-01 grants.

## Author contributions

ISK, ZLG, KSP and BGB conceived the study. ZLG generated atrial and ventricular Hi-C 3.0 datasets, bulk atrial CM RNA-seq and TBX5 ChIP-seq datasets and all snm3C-seq samples. ZLG and AJH generated TBX5 allelic series Hi-C 3.0 datasets. ZLG generated CTCF ChIP-seq datasets with assistance and optimization from CJ. SK processed and analyzed all Hi-C 3.0 data, except dcHiC, which was analyzed by SZ. ZLG processed ChIP-seq and bulk RNA-seq data, which were further analyzed by SK. SZ analyzed snm3C-seq datasets. KSR analyzed scRNA-seq datasets for TBX5 allelic series with supervision and input from ISK. VK generated and analyzed bulk ventricular CM RNA-seq data. PKL, KD, BY and WMB generated and processed snm3C-seq data with supervision and input from NRZ and BR. ZLG, SK, SZ, ISK, KSP and BGB interpreted the data. ZLG, SK and SZ prepared figures. ZLG, SK, SZ, and NRZ wrote the original draft manuscript. ZLG, SK, SZ, ISK, KSP and BGB edited the manuscript with contributions from all co-authors. All authors approved the manuscript.

## Data and materials availability

Cell lines are available under a material transfer agreement. Original Hi-C 3.0, bulk RNA-seq, and TBX5 and CTCF ChIP-seq data are available under NCBI BioProject PRJNA1199549. Original snm3c-seq data are available under NCBI BioProject PRJNA1200909.

