## Supplemental Figures for "Dose-dependent sensitivity of human 3D chromatin to a heart disease-linked transcription factor"

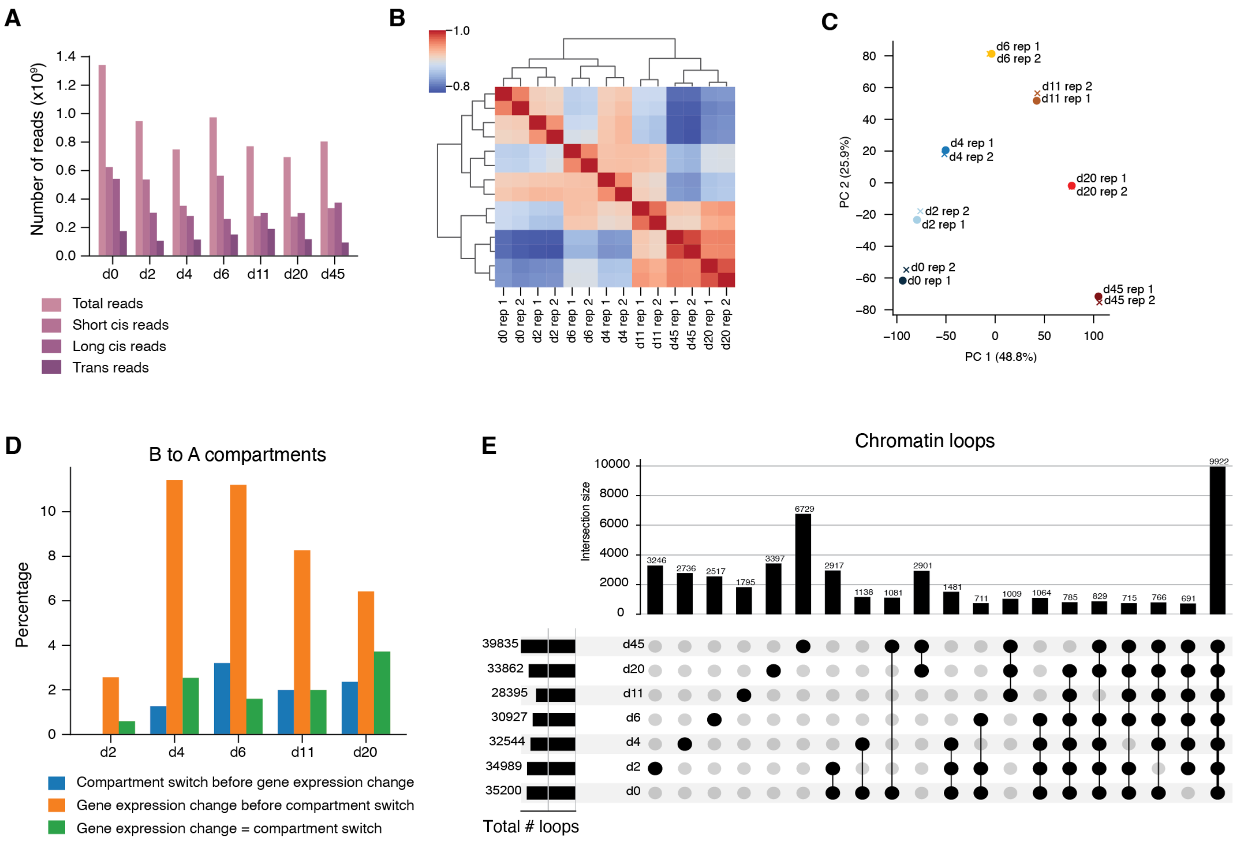


**Fig. S1. Quality control data for atrial cardiac Hi-C 3.0 libraries.** (**A**) Total cis and trans reads per time point. (**B**) Heatmap and (**C**) principal component plot of samples used for atrial differentiation Hi-C 3.0 libraries. (**D**) Proportion of genes with increased expression during atrial differentiation whose expression change precedes, equals or follows a compartment switch from B to A. X-axis indicates the day of differentiation that the genes were increased in expression. (**E**) Upset plot of chromatin loops, indicating consensus loops between two or more time points that are gained across atrial CM differentiation. Two biological replicates per time point were merged for analysis and referred to in this figure.

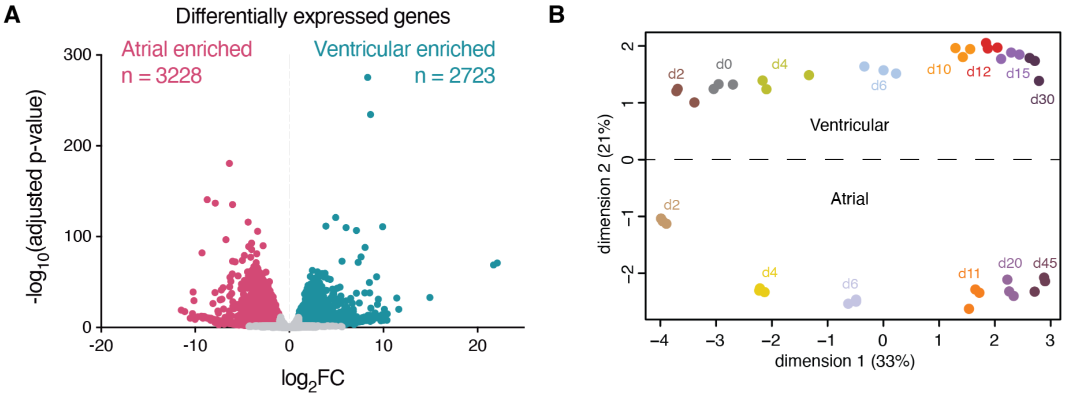


**Fig. S2. Transcriptional differences between atrial and ventricular CMs.** (**A**) Volcano plot of differentially expressed genes between d45 atrial and d30 ventricular CMs. Fuchsia dots are atrial-enriched with $\log_2 FC \leq -1$, $p<0.01$. Teal dots are ventricular-enriched with $\log_2 FC \geq 1$, $p<0.01$. (**B**) Multidimensional scaling plot of bulk RNA-seq samples across ventricular and atrial differentiation. RNA-seq data is n = 3 per time point.

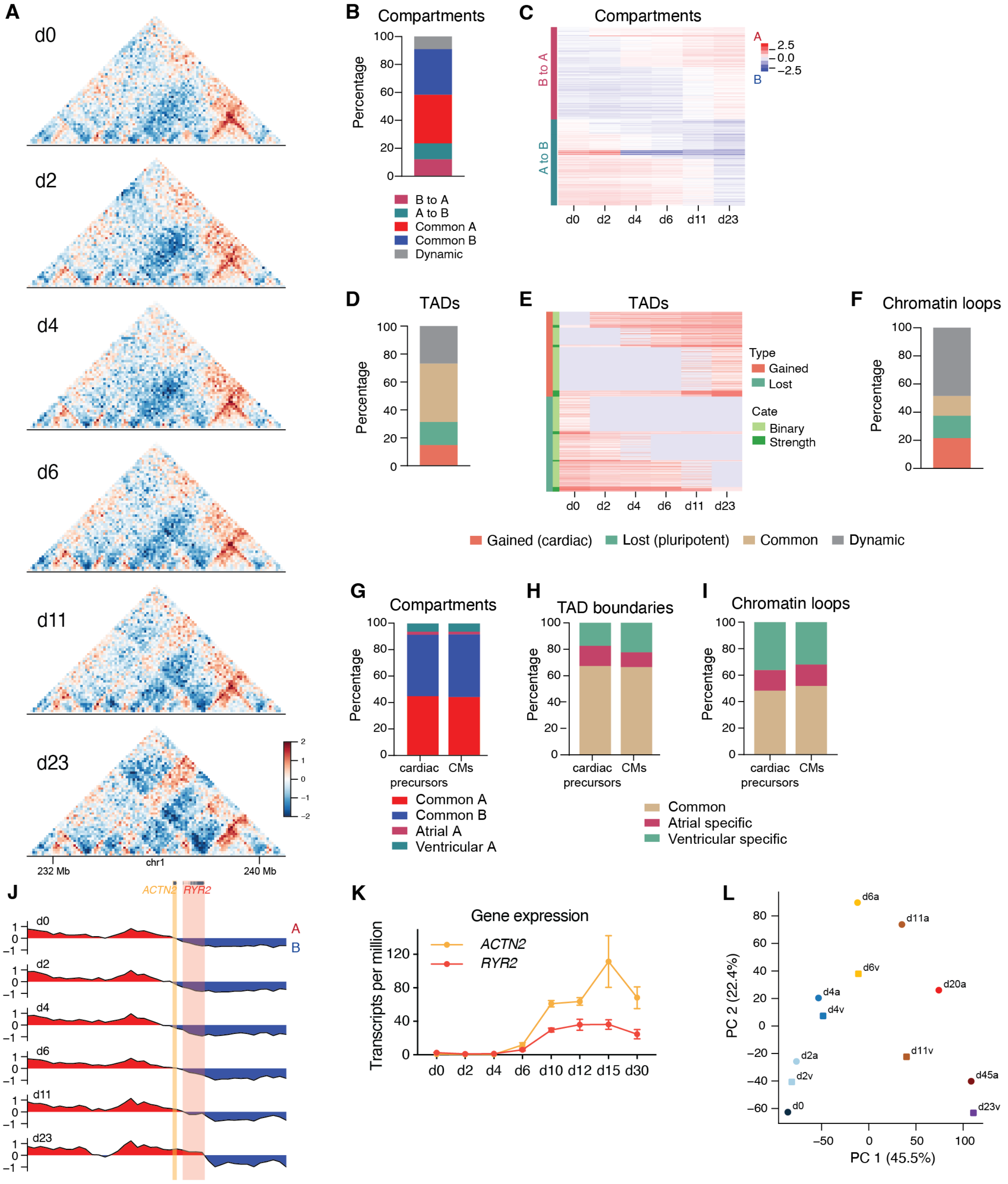


**Fig. S3. Dynamic reorganization of human cardiac chromatin in ventricular differentiation.** (**A**) Observed/expected contact maps in a 10 Mb region around *ACTN2* and *RYR2* in ventricular cardiac differentiation. In contact maps, red indicates regions that are enriched in contacts above expected, blue indicates regions that are depleted. (**B**) Percentage of compartments that switch or remain the same. Dynamic/others indicates compartments that switch more than once. (**C**) Heatmap of compartments that switch from A to B or B to A. (**D**) Percentage of TAD boundaries that are common or changed. (**E**) Heatmap of TADs that change by time point. (**F**) Percentage of chromatin loops that are common or change. (**G-I)** Percentage of compartments, TAD boundaries (F) and loops (G) that are common or differ between ventricular and atrial cardiac precursors (d6), and cardiomyocytes (d23, d45). (**J**) Principal component scores highlighting A/B compartments with *ACTN2* and *RYR2* gene loci highlighted. (**K**) *ACTN2* and *RYR2* gene expression from RNA-seq, mean ± S.E.M. (**L**) Principal component plot of compartment scores of atrial (a) and ventricular (v) Hi-C 3.0 data. Two biological replicates per time point were merged for analysis (except n = 1 for d2 and d4) and referred to in this figure.

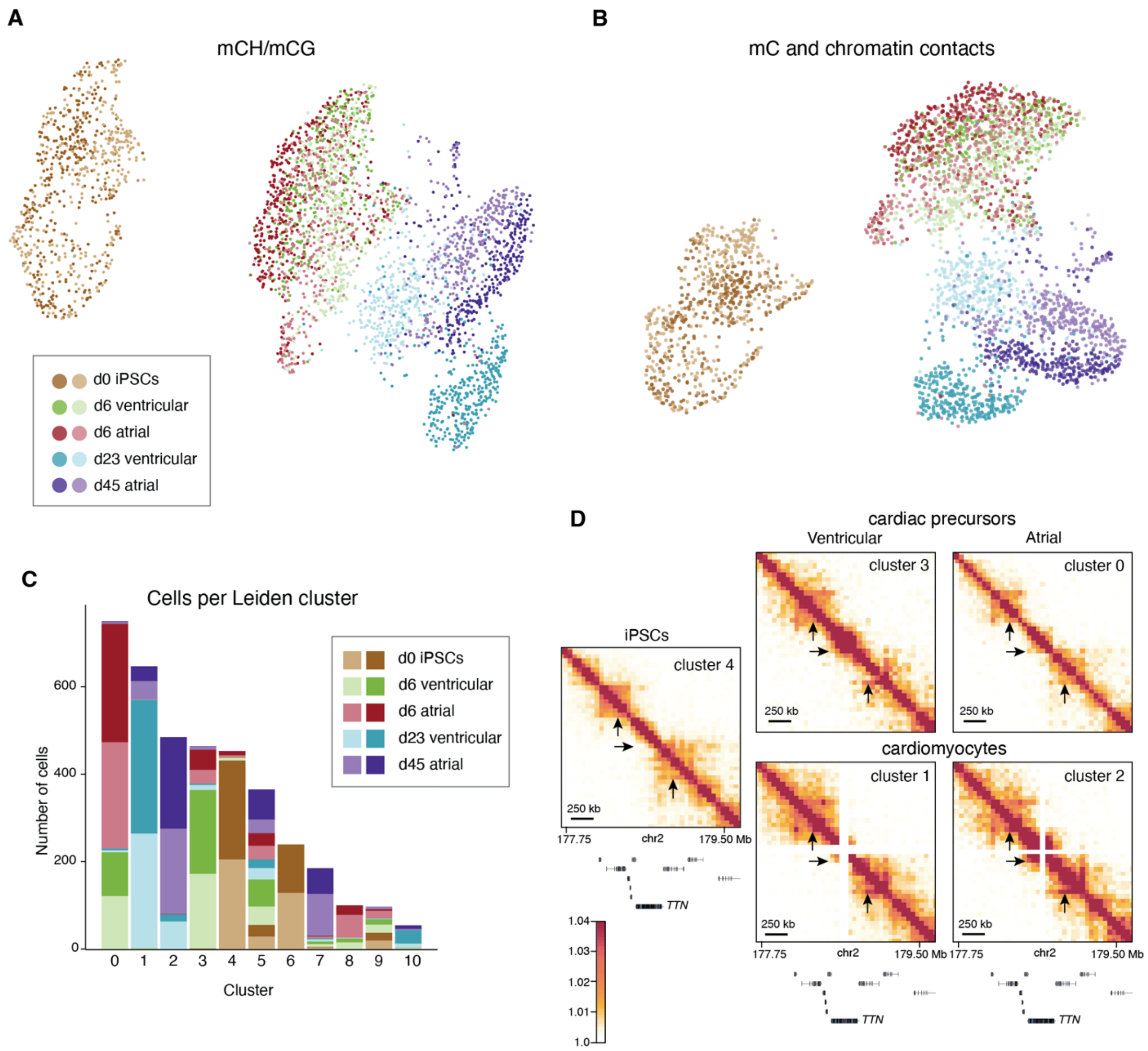


**Fig. S4. snm3c-seq methylation clustering of atrial and ventricular differentiation.** (**A**) UMAP of atrial and ventricular differentiations, where the features are derived from 100 kb genomic regions containing the most variability in mCH and mCG sites across single cells and (**B**) integrating methylation and Fast-Higashi embeddings of Hi-C contacts at 1 Mb resolution. Cells are colored by time point and cell type, with the two shades indicating biological replicates. (**C**) Total number of cells per Leiden cluster colored by time point and cell type. (**D**) Imputed contact matrices of Leiden clusters 0-4 around cardiac enriched gene *TTN*. Black arrows indicate regions of increased contact frequency in cardiomyocyte clusters (1 and 2). Scale shows $\log_2(\text{value}+1)$, where value = balanced contact frequency. Two biological replicates per time point and differentiation were analyzed and referred to in this figure.

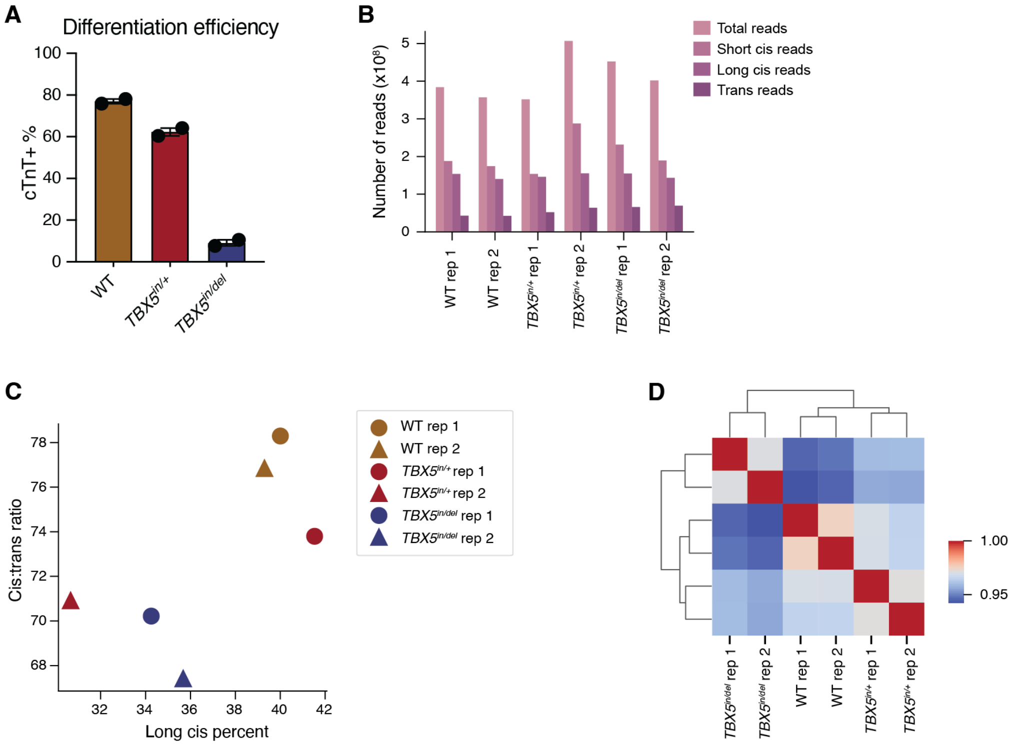


**Fig. S5. Quality control data for *TBX5* allelic series Hi-C 3.0 libraries.** (**A**) Differentiation efficiency as assessed by percentage of cells expressing cTnT per genotype. Mean ± SEM. (**B**) Total cis and trans reads per sample. (**C**) Cis:trans ratio plotted against long cis percent and (**D**) heatmap of samples used for *TBX5* allelic series Hi-C 3.0 libraries. Two biological replicates per genotype were merged for analysis and referred to in this figure.

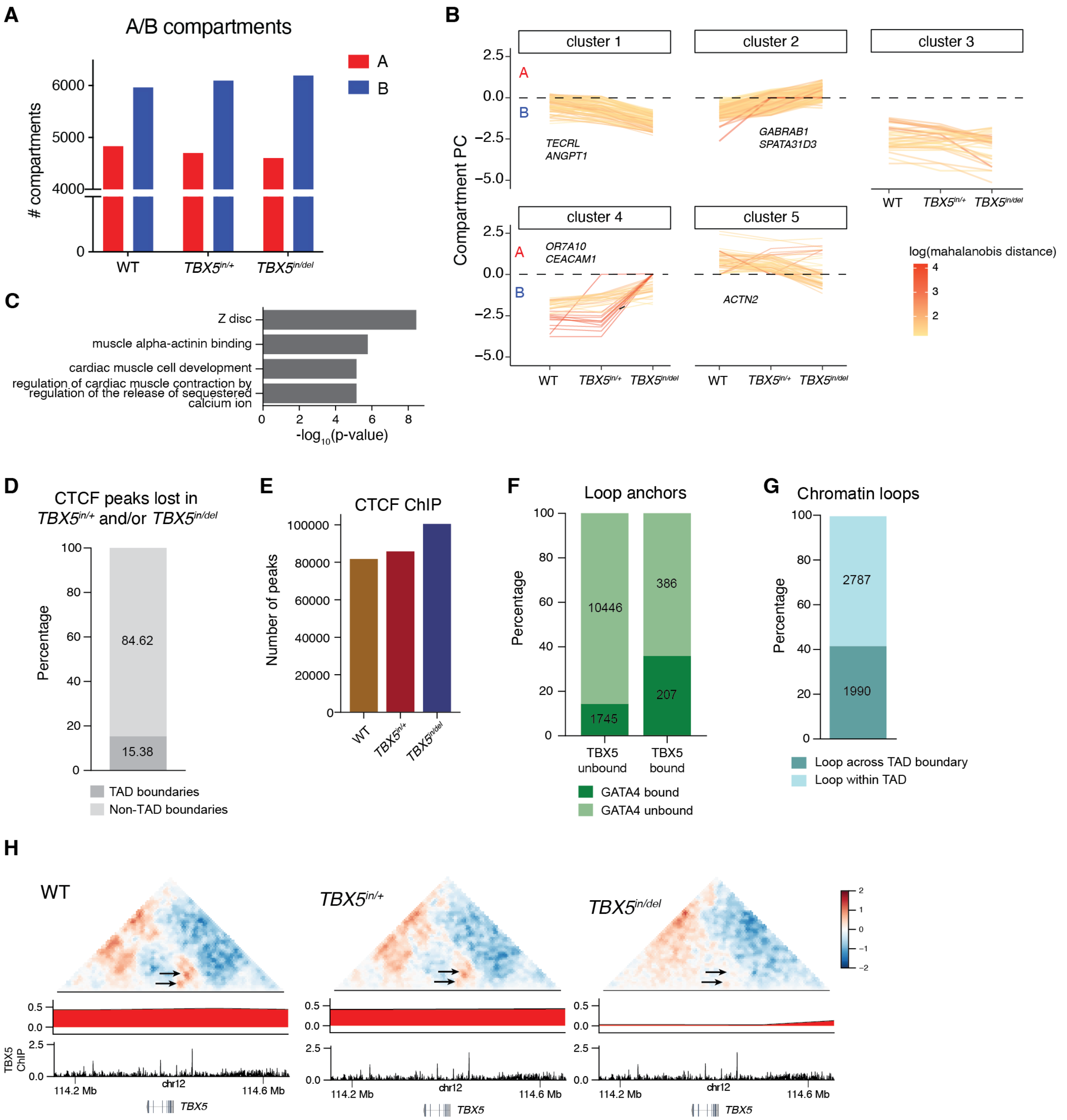


**Fig. S6. Changes in compartments, TADs and chromatin loops in TBX5 allelic series.** (**A**) Total number of A and B compartments per genotype. (**B**) dcHiC clustering of compartment strength. Log-transformed Mahalanobis distance scale showing the degree to which changes in one compartment's score across genotypes is an outlier compared to all compartments. (**C**) Gene Ontology analysis of downregulated genes that are located within compartments switching from A to B in *TBX5*$^{in/+}$ and/or *TBX5*$^{in/del}$ cells. (**D**) Proportion CTCF binding sites that are lost in *TBX5*$^{in/+}$ and/or *TBX5*$^{in/del}$ cells that are associated with WT TAD boundaries. (**E**) Total number of CTCF ChIP peaks detected per genotype. (**F**) Proportion of TBX5 bound or unbound loop anchors by GATA4 co-binding status. (**G**) Proportion of chromatin loops that cross a TAD boundary.

(**H**) Observed/expected contact map with A/B compartment principal components and TBX5 ChIP track around *TBX5*. Arrows indicate reduced contact frequency in $TBX5^{in/+}$ and/or $TBX5^{in/del}$ cells. Two biological replicates per genotype were merged for analysis and referred to in this figure.

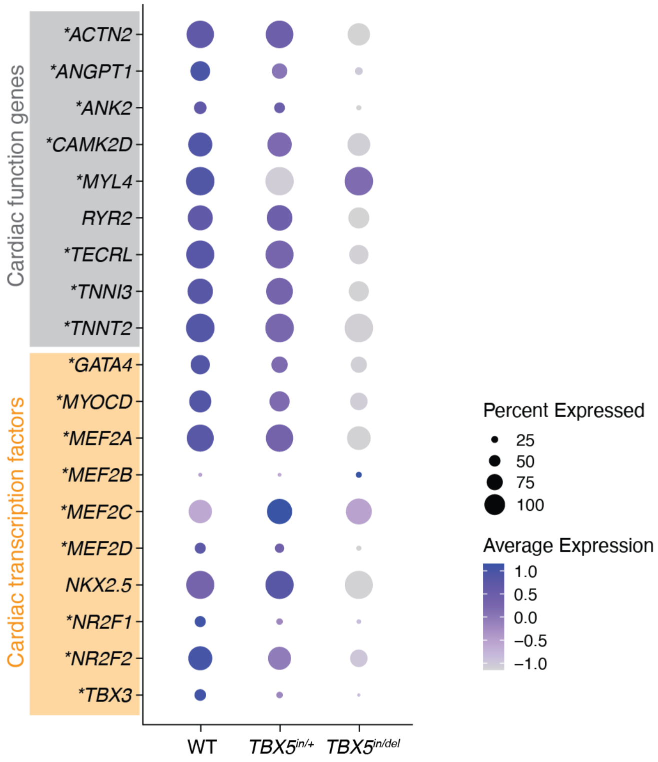


**Fig. S7. Differential gene expression associated with changes in 3D chromatin in TBX5 allelic series.** scRNA-seq clusters enriched for WT, $TBX5^{in/+}$ or $TBX5^{in/del}$ cells were analyzed for differential gene expression. Genes shown are a subset of those associated with altered 3D chromatin in $TBX5^{in/+}$ and/or $TBX5^{in/del}$ cells. Dot size represents the proportion of cells expressing that gene in each cluster. *p<0.01. Two biological replicates per genotype.

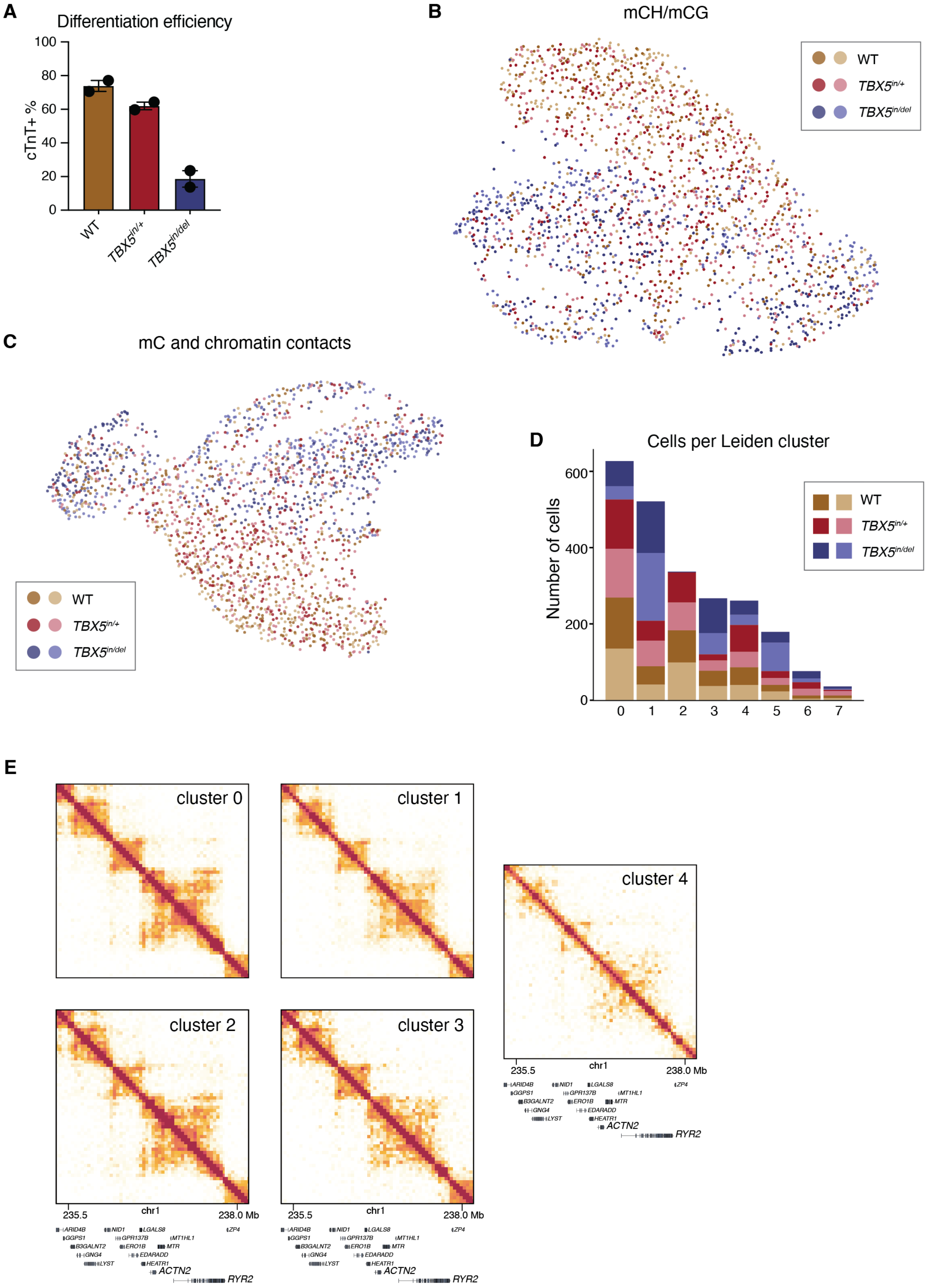


**Fig. S8. snm3c-seq methylation clustering of TBX5 allelic series.** (**A**) Differentiation efficiency as assessed by percentage of cells expressing cTnT per genotype. Mean ±

SEM. (**B**) UMAP of TBX5 allelic series where the features are derived from 100 kb genomic regions containing the most variability in mCH and mCG sites across single cells or (**C**) integrating methylation and Fast-Higashi embeddings of Hi-C contacts at 1 Mb resolution. Cells are colored by genotype, with the two shades indicating biological replicates. (**D**) Total number of cells per Leiden cluster colored by genotype. (**E**) Imputed contact matrices of Leiden clusters 0 to 4 around cardiac enriched genes *ACTN2* and *RYR2*. Scale shows $\log_2$(value+1), where value = balanced contact frequency. Two biological replicates per genotype were analyzed and referred to in this figure.

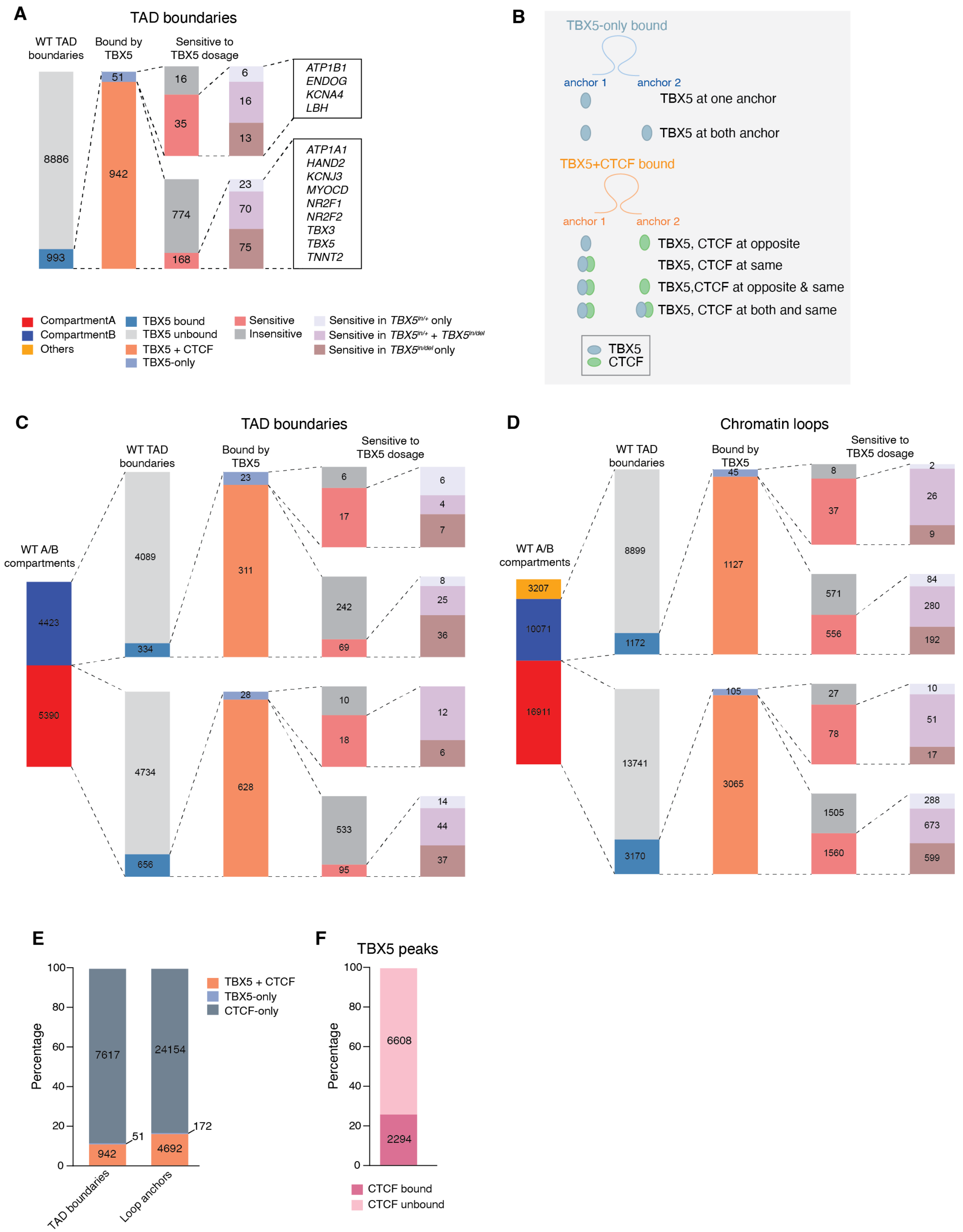


**Fig. S9. TBX5 and CTCF binding at TAD boundaries and chromatin loop anchors for compartment A and B.** (**A**) Percentage of TAD boundaries bound by TBX5

segregated into (TBX5-only or CTCF and TBX5 co-occupied boundaries and proportion of those that lost in *TBX5*$^{in/+}$ and/or *TBX5*$^{in/del}$ samples. Gene list indicates those near sensitive contacts. (**B**) Schematic of classifying TBX5-only and TBX5 + CTCF co-bound loop anchors. (**C**) Percentage of TAD boundaries and (**D**) loop anchors in A and B compartments that are bound by TBX5 segregated into (TBX5-only or CTCF and TBX5 co-occupied boundaries and proportion of those that lost in *TBX5*$^{in/+}$ and/or *TBX5*$^{in/del}$ samples. (**E**) Proportion of TAD boundaries and loop anchors by TBX5 and CTCF binding status. (**F**) Proportion of TBX5 ChIP peaks that overlap with CTCF binding sites. Two biological replicates per genotype were merged for analysis and referred to in this figure.

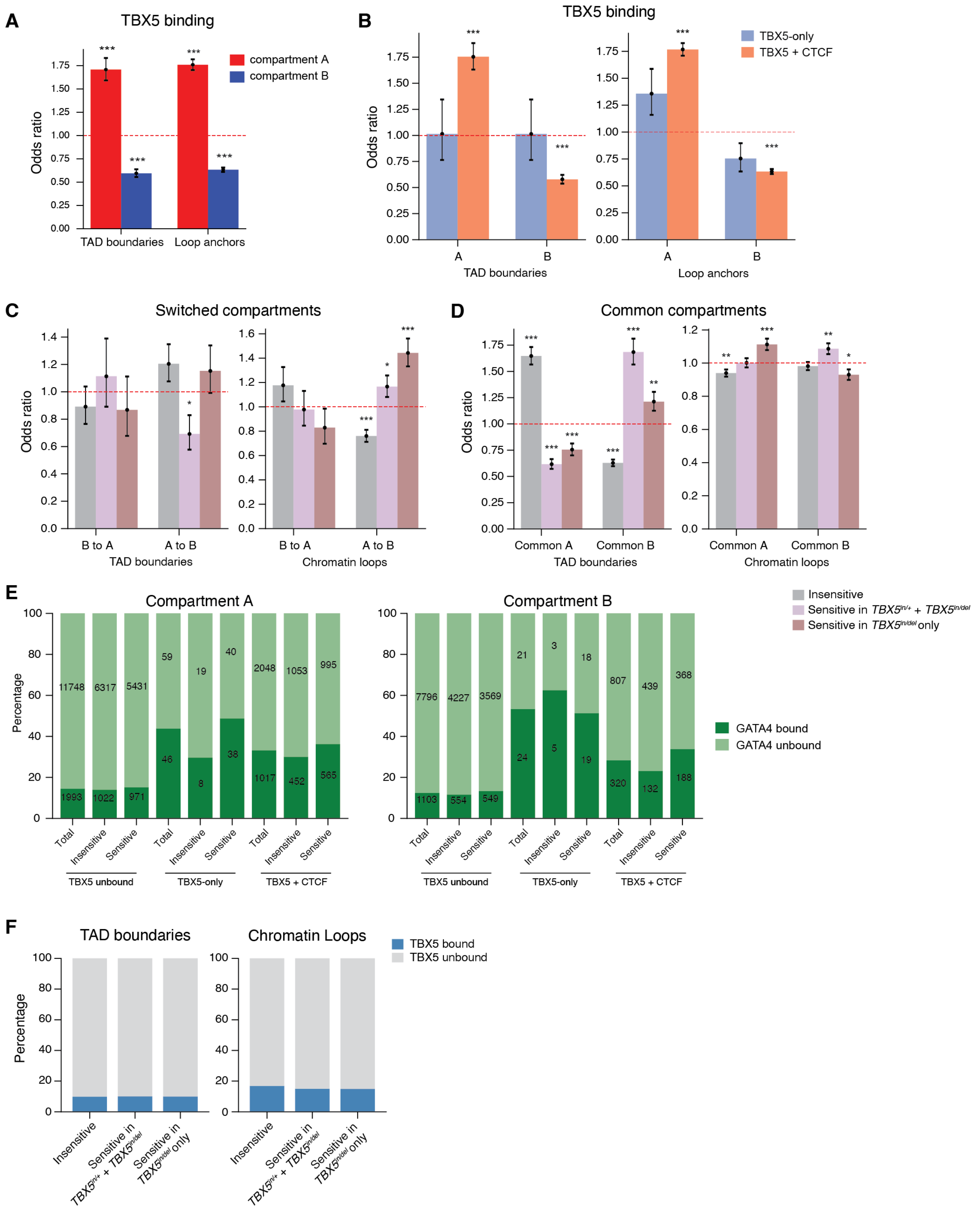


**Fig. S10. Association of TBX5 binding and sensitive regions with compartments.** (**A, B**) Odds ratio of TBX5 binding at TAD boundaries and loop anchors in compartment A and B (A) and separated by TBX5 binding status (B). (**C, D**) Odds ratio of association of TAD boundaries and chromatin loop anchors separated by sensitivity to loss of TBX5 in switched (C) and common compartments (D). (**E**) Proportion of chromatin loops in compartment A and B that are bound by GATA4 separated by sensitivity and TBX5 binding status. (**F**) Proportion of TAD boundaries and chromatin loop anchors that are bound by TBX5 by sensitivity to TBX5 loss. Two biological replicates per genotype were merged for analysis and referred to in this figure. All odds ratios are calculated as each subset against all others not in that subset. Dotted lines at OR = 1 indicates no enrichment. *p<0.05, **p-value<0.01, ***p<0.001, ****p<0.0001.
